# Nde1 Promotes Lis1 Binding to Full-Length Autoinhibited Human Dynein-1

**DOI:** 10.1101/2024.12.30.630764

**Authors:** Jun Yang, Yuanchang Zhao, Pengxin Chai, Ahmet Yildiz, Kai Zhang

## Abstract

Cytoplasmic dynein-1 (dynein) is the primary motor for the retrograde transport of intracellular cargoes along microtubules. The activation of the dynein transport machinery requires the opening of its autoinhibited Phi conformation by Lis1 and Nde1/Ndel1, but the underlying mechanism remains unclear. Using biochemical reconstitution and cryo-electron microscopy, we show that Nde1 significantly enhances Lis1 binding to autoinhibited dynein and facilitates the opening of Phi. We discover a key intermediate step in the dynein activation pathway where a single Lis1 dimer binds between the Phi-like (Phi^L^) motor rings of dynein. In this “Phi^L^-Lis1”, Lis1 interacts with one of the motor domains through its canonical interaction sites at the AAA+ ring and stalk and binds to the newly identified AAA5, AAA6, and linker regions of the other motor domain. Mutagenesis and motility assays confirm the critical role of the Phi^L^-Lis1 interface. This intermediate state is instantly and efficiently formed in the presence of Nde1, but Nde1 is not part of the Phi^L^-Lis1. These findings provide key insights into the mechanism of how Nde1 promotes the Lis1-mediated opening of Phi dynein.

## Introduction

Cytoplasmic dynein is a motor protein essential for transporting nearly all intracellular cargoes toward the minus end of microtubules (MTs) in most eukaryotes. Dynein’s cargoes include organelles, proteins, RNA, viruses, and vesicles^1^. Additionally, dynein plays key roles in other cellular processes such as mitosis and organelle positioning^1,2^. Mutations affecting dynein or its regulatory proteins have been linked to neurodevelopmental and neurodegenerative diseases, including spinal muscular atrophy (SMA), ALS, Huntington’s disease, lissencephaly, and microcephaly^3–9^.

The 1.4 MDa dynein complex contains pairs of six subunits. The largest subunit is the dynein heavy chain (DHC), which contains the N-terminal tail domain and C-terminal motor domain. The tail domain facilitates dimer formation, recruits the dimers of intermediate chain (DIC), light intermediate chain (DLIC), and interacts with dynactin, cargo adaptors and other regulatory proteins^10–13^. The N-terminus of DIC (DIC-N) recruits three pairs of light chains, Robl, LC8, and Tctex^1,14,15^. DIC segments binding to LC8 and Tctex, known as IC-LC tower, play a crucial role in assembling the dynein-dynactin-adaptor (DDA) complex^11^. The dynein motor domain belongs to AAA+ (ATPases Associated with diverse cellular Activities) protein families and consists of six AAA subdomains (AAA1-6)^16,17^. AAA1 is the primary subdomain that hydrolyzes ATP to power dynein motility along MTs. The dynein motor domain attaches to MTs via the coiled-coil stalk and the MT-binding domain (MTBD) and connects to the tail through the linker domain located at the surface of the AAA+ ring^17–20^.

Dynein alternates between two key conformational states: the autoinhibited Phi (similar to the Greek letter “φ”) and the open conformations^14,21^. Phi dynein adopts a compact structure that limits its interactions with MTs and dynactin, which serves to minimize unnecessary ATP hydrolysis when motor protein is not engaged in active transport^14,22^. Opening of the Phi enables dynein to assemble with dynactin and the cargo adaptor, allowing processive movement along MTs^10,14^, a process supported by dynein regulators Nde1/Ndel1 and Lis1^3,15,23–30^. However, the underlying mechanism promoting Phi to open transition is unclear.

Lis1, the first gene identified in relation to a neuronal migration disease, plays a crucial role in dynein-related function^3,11,31–39^. Lis1 possesses an N-terminal LisH domain that facilitates dimerization and a C-terminal WD-40 β-propeller domain that binds to Nde1/Ndel1, dynein, and other proteins^3^. Both domains are important at different stages of dynein activation^40–42^. The LisH domain interacts with dynactin p150 and DIC-N, thus promoting the recruitment of dynactin and adaptors at the later stage of dynein activation^11^. WD-40 domains of Lis1 can directly bind to the dynein motor domain at the AAA3-AAA4 sites (site-ring) and stalk coiled-coil (site-stalk)^11,38,43^. Functional studies proposed that Lis1 facilitates the formation of highly processive DDA complex by favoring the release of Phi dynein and stabilizing open dynein^34–37,44^. Lis1 binding is thought to be incompatible with Phi dynein based on the steric clash when docking a Lis1 β-propeller to the motor domains of Phi dynein^3,35,37,44^. However, there is no direct biochemical or structural evidence of whether Lis1 can open Phi dynein and intermediary states that facilitate opening of Phi dynein remain unclear. A recent structural study on tail-truncated yeast dynein motor domains reported a “Chi” conformation, in which two Lis1 dimers are wedged between two AAA+ rings^37^, suggesting that two Lis1 dimers between the motor domains are required to crack open Phi dynein. Furthermore, it is well-known that the AAA+ ring undergoes substantial conformational changes under different nucleotide-binding states, which can potentially regulate dynein–Lis1 binding. A more recent work has demonstrated that specific nucleotide “codes” at the three variable nucleotide-binding sites (AAA1, 3, and 4) govern the stoichiometry of dynein–Lis1 interactions by tuning their binding affinity at two distinct locations^45^. However, the intermediate structure for full-length human dynein alone bound to Lis1 is still lacking and whether dynein forms a Chi conformation at the initial state of its activation remains unclear.

Nde1 and its paralog Ndel1^12,23^, are critical for all dynein-mediated functions in cell division^28,46^, cargo trafficking^47,48^ and neuronal migration^49,50^. Nde1/Ndel1 is predicted to be composed of the N-terminal coiled-coil domain and the C-terminal unstructured region^51–53^. The coiled-coil domain interacts with DIC-N, which overlaps with the DIC binding site of the p150 subunit of dynactin^52,54,55^. It also interacts with the WD-40 domains of Lis1, and its binding site on WD-40 domain overlaps with the Lis1 binding site of dynein^26,52^. The multi-protein interaction modes of Nde1/Ndel1, along with its overlapping binding sites with other proteins, make the functional interpretation of Nde1/Ndel1 elusive. It has been proposed that Nde1 tethers Lis1 to dynein^12,26,56–58^ and promotes Lis1-mediated activation of dynein^11,26^ (**Fig. 1a**). According to this model, Phi dynein first adopts an ‘open’ conformation. Lis1 stabilizes the open dynein by preventing it from transitioning back to Phi, thus favoring the assembly of the DDA complex^11,34,35,44^. Consistent with this model, overexpression of Lis1 can rescue the deletion of Nde1/Ndel1 in cells^58,59^ and Nde1/Ndel1-mediated recruitment of Lis1 to dynein enhances DDA assembly in vitro^26^. An alternative model suggests that Ndel1 negatively regulates dynein activation by competing with p150 for DIC-N binding and by sequestering Lis1 away from dynein^52^. Consequently, how Lis1 and Nde1/Ndel1 form a complex with dynein and promote the opening of Phi dynein is not well understood.

**Fig. 1.**
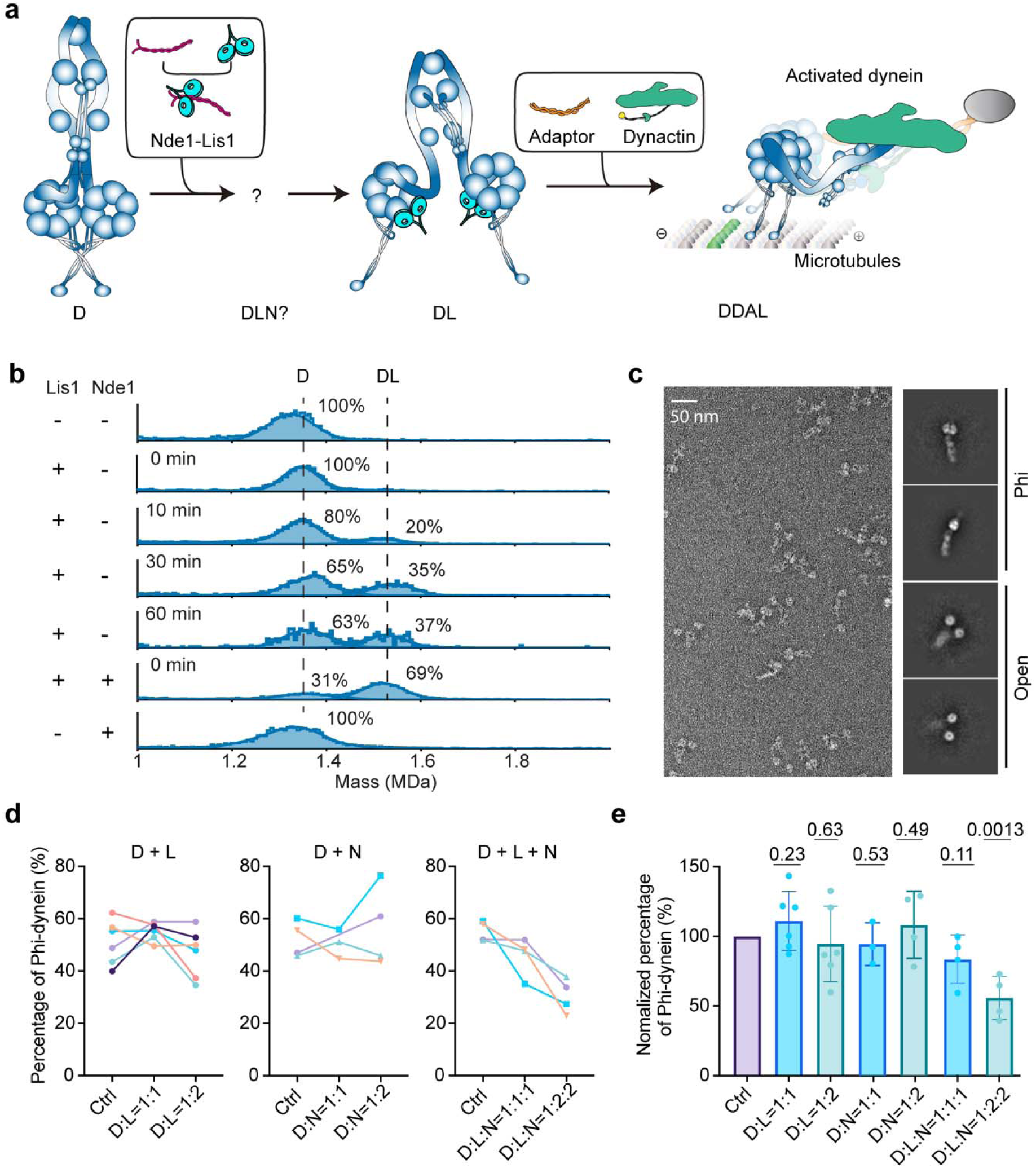
Nde1 promotes Lis1 binding to Phi dynein and cooperatively releases dynein autoinhibition. **a**, Schematic of dynein activation. Nde1 tethers Lis1 to dynein and promotes Lis1-mediated formation of the active dynein-dynactin-adaptor-Lis1 (DDAL) complex. Intermediate states between the association of Lis1-Nde1 and opening of dynein are unknown. **b**, MP shows that Lis1 alone slowly binds to dynein in tens of minutes, whereas Nde1 promotes more rapid and efficient binding of Lis1 to dynein. Dynein, Lis1 and Nde1 were included at a 1:2:2 ratio. Only one Lis1 dimer is tethered to one dynein. Solid curves represent a fit to multiple Gaussians to predict the average mass and percentage of each population. **c**, A representative image of dynein motors captured using negative stain electron microscopy. Without Lis1 and Nde1, dynein is distributed almost equally between Phi and open conformations. The percentage of Phi was quantified after incubating dynein with Lis1, Nde1, or both proteins for 90 min. The percentage of Phi (**d**) and the relative change of Phi (**e**) in the presence and absence of Lis1 and Nde1 (mean ± s.d.; from left to right, n=14, 6, 6, 3, 4, 4, 4 from three or more independent experiments). P values are calculated from a two-tailed *t* test. The control (Ctrl) of panel e represents dynein alone from the three groups of panel d (Ctrl) normalized to 100%.

To understand the mechanism by which Nde1/Ndel1 and Lis1 prime dynein for the DDA assembly, we investigated how human Lis1 and Nde1 affect the conformational states of full-length human dynein using biochemical reconstitution and electron microscopy. We showed that Nde1 promotes the formation of the dynein-Lis1 complex. Using negative-stain EM, we provide direct evidence that Lis1 or Nde1 alone has little effect on the equilibrium between the Phi and open conformations of dynein, but Lis1 and Nde1 together significantly bias the equilibrium toward the open conformation, demonstrating that Nde1 acts like a molecular chaperone to promote Lis1-mediated opening of dynein. Cryo-EM imaging of dynein-Lis1 in the presence or absence of Nde1 captures a new intermediate and rate-limiting state during the dynein activation process, characterized by a single Lis1 dimer binding between the motor domains of Phi dynein. Lis1 binding to Phi dynein causes the rotation between two motor domains to relieve the steric clash, forming a Phi-like dynein and Lis1 complex, named “Phi^L^-Lis1”. While Lis1 binds to one of the motor domains through its canonical interaction sites, it also forms new interfaces between the other motor domain at the AAA5, AAA6, and linker regions. Mutagenesis at the novel interfaces together with single molecule motility assays supported the critical role of this Phi^L^-Lis1 during the dynein activation process. Collectively, our findings shed light on the roles of Nde1 and Lis1 in the dynein activation pathway.

## Results

### Nde1 promotes Lis1 binding to Phi dynein and cooperatively releases dynein autoinhibition

To determine whether Nde1 promotes Lis1 binding to dynein, we performed mass photometry (MP) assays to assess the Lis1 and Nde1 binding to full-length human dynein under different conditions. In the absence of Nde1, Lis1 alone exhibited an increased binding to dynein in a time-dependent manner, with the formation of a 37% 1:1 dynein:Lis1 (DL) complex within 60 minutes. The inclusion of Nde1 significantly enhanced the dynein-Lis1 binding (69% of complex formation) in less than a minute, consistent with Nde1/Ndel1-mediated tethering of Lis1 to dynein^12,26,57^ (**Fig. 1b**). To further test if Nde1 preferentially recruits Lis1 to Phi or open conformation, we performed these assays using the Phi mutant of dynein^14^ that only forms open conformation. Lis1 was readily bound to open dynein, but the addition of Nde1 did not further enhance Lis1 binding to dynein (**Extended Data Fig.1**), suggesting that Nde1 is required for Lis1 recruitment to Phi dynein. These results are also consistent with the previous observation that the Nde1 addition does not further enhance Lis1-mediated activation of DDA complexes assembled with the Phi dynein mutant^26^. Interestingly, the formation of the dynein-Lis1-Nde1 tripartite complex was not observed (**Fig. 1b and Extended Data Fig.1**), regardless of different nucleotide conditions (**Extended Data Fig. 2**). Although previous reports indicated that Nde1 can interact with DIC-N in single molecule imaging^12,26^ and pull-down assays^26,52^, we also did not detect the dynein-Nde1 complex (**Fig. 1b**, and **Extended Data Fig. 2**), suggesting that Nde1 rapidly dissociates from the complex after handing off Lis1 to dynein^12^.

To determine whether Nde1-mediated Lis1 recruitment to dynein shifts the equilibrium between Phi and open conformations, we used the negative stain EM imaging^14^ to quantify the ratio of Phi dynein in the presence and absence of Lis1 and Nde1. Specifically, we used freshly prepared dynein with ∼50% of motors forming the Phi and then incubated dynein with Lis1 and Nde1 (**Fig. 1c and Extended Data Fig. 3**) in the presence of ATP. We found that Lis1 or Nde1 alone does not change the Phi ratio compared with the control (**Fig. 1d-e**). However, the Phi ratio decreased 44% when we incubated dynein with both Lis1 and Nde1 at a ratio of 1:2:2 (**Fig. 1d-e**). Collectively, our results demonstrate that Nde1 specifically promotes Lis1 binding to Phi dynein and facilitates opening of this autoinhibited conformation.

### A novel Phi^L^-Lis1 structure

Our MP results suggest that there is a rate-limiting step of Lis1 binding to Phi dynein, and this step can be significantly accelerated by the addition of Nde1 (**Fig. 1b**). We used cryo-EM to capture the potential intermediate states to reveal the structural basis of this process. We focused on the particles that form the autoinhibited dynein (**Fig. 2a, b and Extended Data Fig. 4, 5**). In the absence of Nde1, we unexpectedly observed a novel structure in which a Lis1 dimer is wedged between the two stacked motor rings of Phi dynein. The dynein in this complex shows a compact conformation, similar to but not the same as the previously reported Phi structure^14^ and we referred this complex as the “Phi^L^-Lis1” (**Fig. 2a, c and Supplementary Video 1**). Despite excess Lis1, nearly half of Phi dynein does not bind to Lis1 (42.7% Phi vs. 57.3% Phi^L^-Lis1) (**Fig. 2a**). Notably, we did not observe the “Chi” (two Lis1s bound to dynein), suggesting that the “Chi” conformation may be specific to yeast dynein or may form when truncated dynein containing only the motor domains is used instead of full-length motor^37^.

**Fig. 2.**
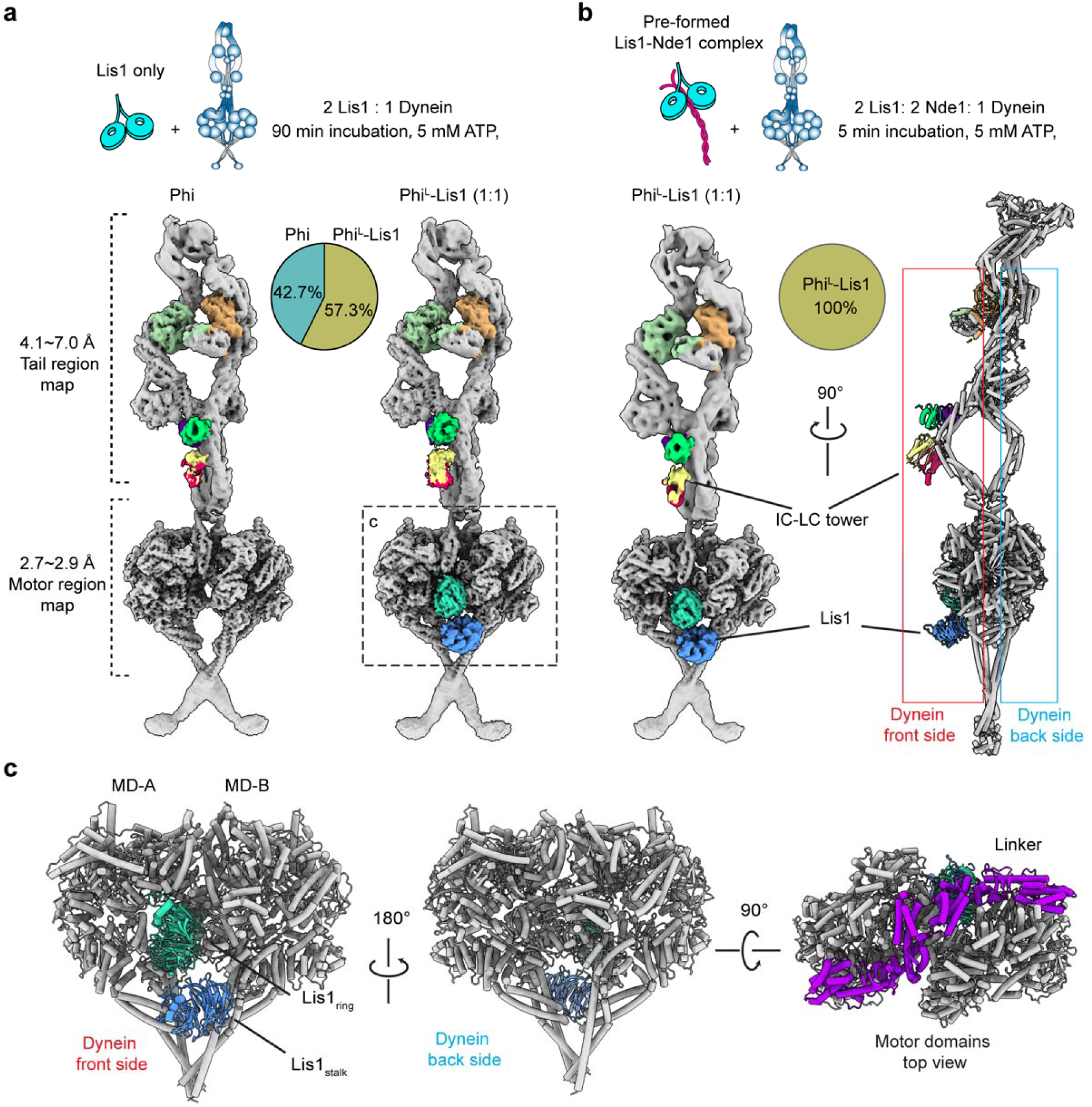
The structure of Phi^L^-Lis1 complex. **a,** (Top) Full length human dynein was incubated with Lis1 with a molar ratio of 1:2 on ice for 90 min before flash frozen. (Bottom) Cryo-EM maps of Phi and Phi^L^-Lis1 complex structures in front view and percentages of particles with these two conformations (open dynein excluded). **b**, (Top) Dynein was incubated with Lis1 and Nde1 with a molar ratio of 1:2:2 on ice for 5 min before flash frozen. (Bottom) The structure of Phi^L^-Lis1 obtained under this condition is shown in front view and side view. All particles were classified into the Phi^L^-Lis1 conformation and Phi conformation could not be detected**. c**, Model of the Phi^L^-Lis1 motor domains is shown in three views. One Lis1 dimer is clamped between MD-A and MD-B of Phi^L^-Lis1 on the front side, while the back side of the motor domains is vacant, with no Lis1 bound. The top view reveals that the linker regions (purple) of MD-A and -B interact with each other.

In the Phi^L^-Lis1 structure, Lis1 appears on the same side as the IC-LC tower of dynein (front side), independent of the presence of Nde1 (**Fig. 2a, b**). Based on this structural observation, along with previous evidence indicating that DIC-N can bind to both Lis1^11^ and Nde1^12^, we speculate that the transient dynein-Lis1-Nde1 forms only on the IC-LC tower side. Although we do not observe clear cryo-EM densities for the DIC-N, it is possible that the DIC-N can weakly interact with Lis1, thus recruiting Lis1 to the front side of dynein.

Previous results suggest Lis1 can affect dynein’s mechanochemical cycle and nucleotide state^11,60,61^. The high-resolution structure of the motor domain enabled us to identify and compare the nucleotide states of the motor domains in Phi and Phi^L^-Lis1 structures. Specifically, we found that AAA1 pockets of both motor domains exhibit the same ADP-Mg^2+^ density (**Extended Data Fig.6a, b**), accompanied by a flexible sensor-I loop, indicating the intermediate state of Pi releasing (**Extended Data Fig. 6c, d**). Similarly, AAA3 pockets show clear ADP binding (**Extended Data Fig.6a, b**), indicating that Lis1 binding does not influence the nucleotide states of the motor domains of autoinhibited dynein.

We next premixed equimolar Nde1 and Lis1, then added dynein to achieve a final 1:2:2 ratio of dynein:Lis1:Nde1 (**Fig. 2b**). Strikingly, we obtained a similar Phi^L^-Lis1 structure, but no longer observed the Phi dynein alone in the presence of Nde1 (**Fig. 2a, b and Supplementary Video 1**). This is consistent with the MP analysis showing that Nde1 instantly promotes Lis1 binding to dynein (**Fig. 1b**) and forming Phi^L^-Lis1. These results suggest that Phi^L^-Lis1 formation is a rate-limiting intermediate state before dynein opening. Remarkably, none of the 3D classes shows clear Nde1 density in Phi^L^-Lis1, suggesting that Nde1 is not part of this complex.

We also analyzed individual motor domains of open dynein and did not observe an increase in the propensity of the dynein-Lis1 complex compared to open dynein alone in the presence of Nde1 (**Extended Data Fig. 7**). Consistent with MP of the Phi dynein mutant^14^ (**Extended Data Fig. 1**), these results show that Nde1 promotes Lis1 binding to Phi dynein to form Phi^L^-Lis1, whereas Lis1 can readily bind to open dynein and does not require Nde1.

### Lis1 induces a relative rotation in the Phi conformation to accommodate its binding

Consistent with previous reports^35,37,44^, our structural analysis reveals a severe steric clash between Lis1 and Phi motor domain A (**Fig. 3a**) and this needs to be relieved to accommodate Lis1’s binding in Phi^L^-Lis1 (**Fig. 3b**). Comparing Phi to Phi^L^-Lis1 reveals a relative rotation between the two motor domains, which results in the groove on the front side of dynein becoming larger than the corresponding groove on the back side (**Fig. 3c, d**). The enlarged groove on the front side allows Lis1 to fit between Phi^L^ motor domains. However, docking of Lis1 to the back side with a smaller groove shows severe clashes, explaining why there is only one Lis1 present in Phi^L^-Lis1 (**Fig. 3e**). The rotation between the motor domains in Phi^L^-Lis1 also causes a slight anticlockwise twist in the neck region (**Fig. 3d**). This twist likely promotes the unwinding of the tail, generating a trend toward an open conformation of dynein. We concluded that Lis1 binding induces a rotation of Phi dynein motor domains to avoid steric clash with Lis1 (**Fig. 3 and Supplementary Video 1**).

**Fig. 3.**
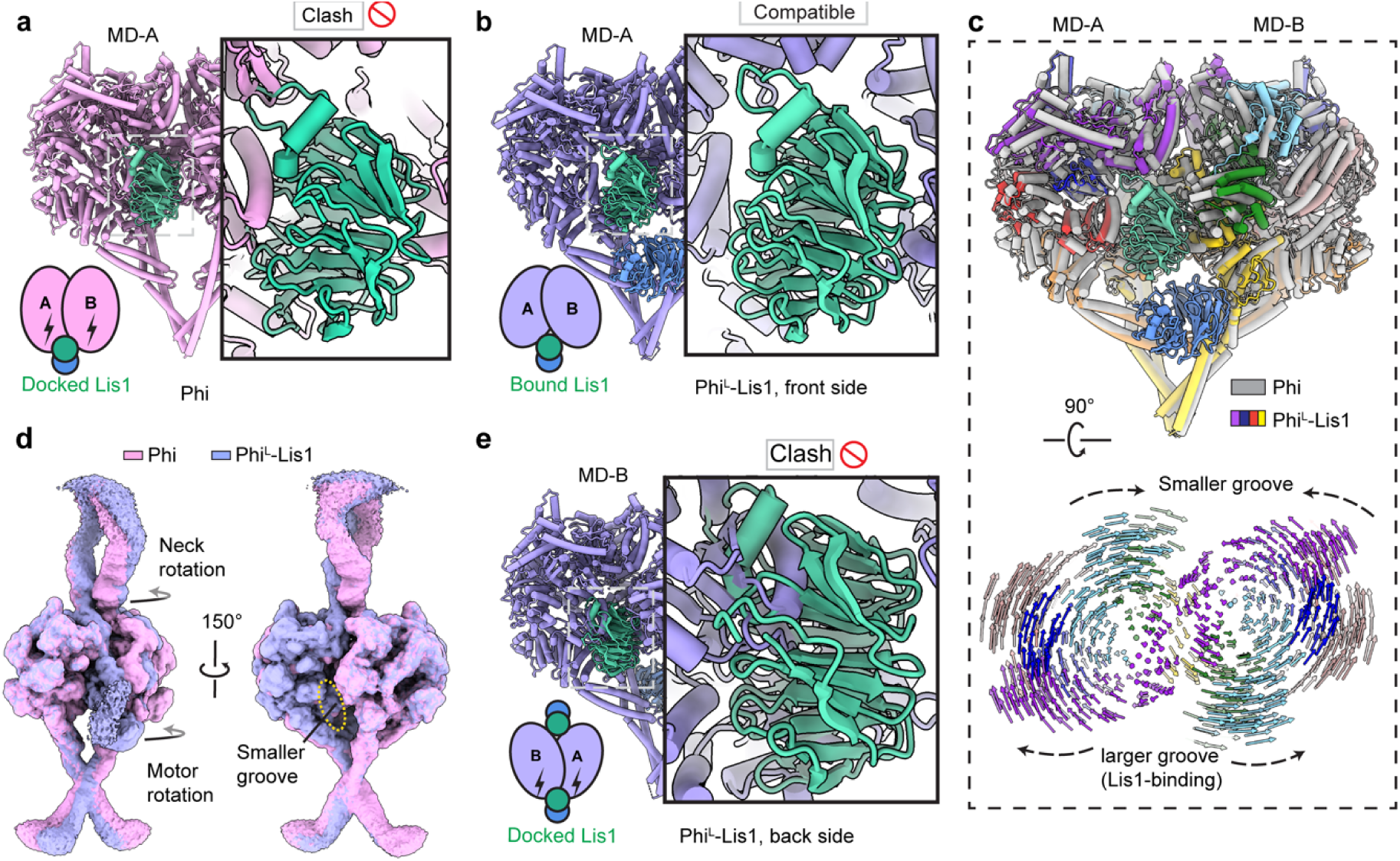
Lis1 induces a transition in the Phi conformation to accommodate its binding. **a**, Docking of a Lis1 monomer between dynein motor rings shows a clash with the rigid canonical Phi MD-A. **b**, A Lis1 dimer bound to dynein motors on the front side is compatible with the coordinated Phi^L^ structure. **c**, (Top) The superimposition of the Phi and Phi^L^-Lis1 motor domains. Phi and Phi^L^-Lis1 are colored with grey and rainbow respectively. (Bottom) Comparison of the motor domains between Phi^L^-Lis1 and canonical Phi. Lis1 is hidden for clarity. Vectors present interatomic distance of pairwise Cα atoms between the Phi to Phi^L^-Lis1 structures. Lis1 binding induces a slight rotation of the dynein motor domains from the Lis1-bound (front) side toward the back, resulting in a larger groove at the front side compared to the smaller groove at the back. **d**, Overlay of the motor domain maps of Phi and Phi^L^-Lis1 with MD-A aligned at a lower contour level. The rotation of Phi^L^-Lis1 motor domain induces rotation in the neck region, showing a slight unwinding trend, which contributes to the formation of a smaller groove on the back side. **e**, Docked Lis1 on the back side of Phi^L^-Lis1 causes a significant clash with MD-B.

### Novel interactions identified in Phi^L^-Lis1

In Phi^L^-Lis1, WD-40 domains of Lis1 interact with both motor domains of dynein (MD-A and MD-B). While Lis1 interacts with MD-B through its canonical interaction sites at the AAA3, AAA4, and AAA5 regions^38,43,62^, we observed previously uncharacterized interaction sites of Lis1 with the linker, AAA6, and AAA5 regions of MD-A (**Fig. 4a, Extended Data Fig. 8 and Supplementary Video 1**). Sequence alignment of Lis1 homologs shows that MD-A and Lis1 interface is highly conserved among higher eukaryotes but less conserved in yeast (**Extended Data Fig. 9**), suggesting different regulatory roles of Lis1 between higher eukaryotes and yeast. Structural comparison indicates that two motor domains in Phi^L^-Lis1 adopt an almost identical conformation. The root-mean-squared-displacement of alpha carbon atoms (Cα-RMSD) of the two motor domains was 0.513 Å (**Fig. 4b**). Additionally, the MD-B bound with Lis1_ring_ and Lis1_stalk_ in our results also shows no significant difference from the structure of Lis1 bound to the human dynein motor domain^38^ (Cα-RMSD: 0.868 Å) (**Fig. 4c**), suggesting Lis1 binding does not induce structural changes within an individual motor domain.

**Fig. 4.**
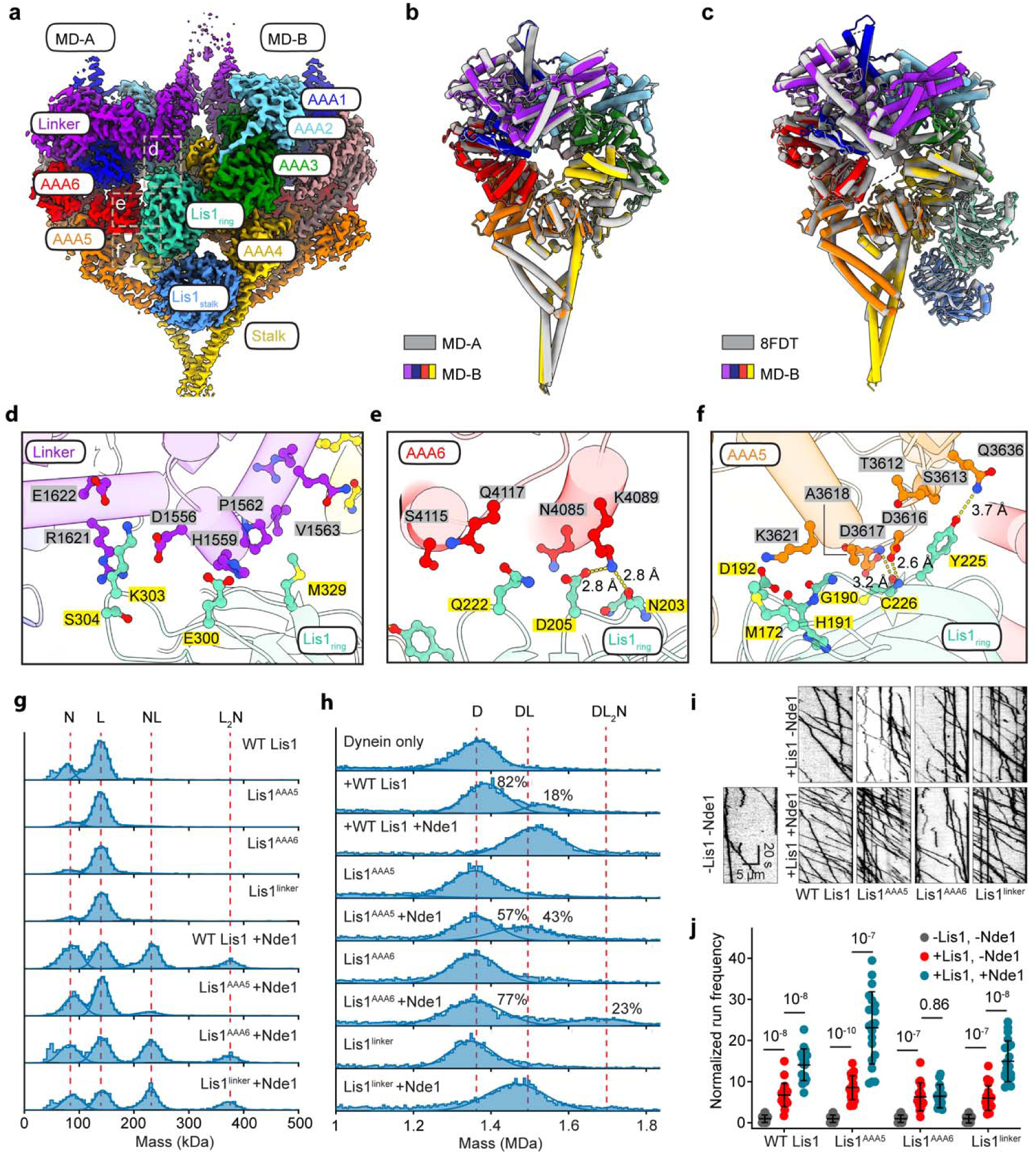
Novel interactions identified between Lis1_ring_ and MD-A in Phi^L^-Lis1. **a**, Cryo-EM density map highlighting the subdomains involved in the interface between Lis1 and the dynein motor. Lis1 and subdomains of MD-A and MD-B are colored separately. Novel interfaces between MD-A and Lis1_ring_ are marked by dash rectangle and enlarged in panels d, e, and f. **b**, Comparison of MD-A (grey) and -B (rainbow) of Phi^L^-Lis1. **c**, Comparison of MD-B (rainbow) with the Lis1-bound structure of the motor domain of human dynein (PDB: 8FDT, grey). Representative interactions located at linker-Lis1_ring_ (**d**), AAA6-Lis1_ring_ (**e**), and AAA5-Lis1_ring_ (**f**) interface are shown with stick mode, colored according to their respective subdomains. **g**, MP profiles illustrate the interaction of Nde1 (N) with WT Lis1 (L) and Lis1 mutants (Lis1^AAA5^, Lis1^AAA6^, and Lis1^linker^). Nde1 interacts with one (NL) or two (NL_2_) Lis1 dimers. Lis1^AAA5^ exhibits a reduced binding percentage to Nde1. **h**. MP shows the binding of Lis1 mutants to dynein with or without Nde1. Dynein, Lis1 and Nde1 were incubated for 2 minutes at 1:2:2 ratio (D: dynein only, DL: one dynein and one Lis1, DL_2_N: one dynein, two Lis1, and one Nde1). **i**, Representative kymographs show the motility of WT DDR complexes with or without Nde1 and Lis1. **j**, Run frequency of WT DDR with or without Nde1 and Lis1 (mean ± s.d.; n = 20 MTs for each condition; statistics from two independent experiments). Results were normalized to the -Lis1, -Nde1 condition.

The interactions between MD-A and Lis1_ring_ are notably compact (**Fig. 4a and Extended Data Fig. 8**). The WD-40 domain of Lis1 interacts with dynein MD-A at regions distributed across the linker, AAA6, and AAA5 region (**Fig. 4d-f**). Specifically, at the linker-Lis1 binding site, the side chains of M329 and E300 of Lis1 engage in hydrophobic and polar interactions with the side chains of V1563, P1562 and H1559 of dynein. Additionally, K303 and S304, located on the flexible loop of the Lis1 WD-40 surface, interact with R1621, D1556, and E1622 of linker region (**Fig. 4d, Extended Data Fig. 8a**).

Within the AAA6 interaction region, the interface is characterized by polar interactions involving N203, D205, and Q222 of the Lis1 WD-40 domain and K4089, N4085, Q4117, and S4115 of MD-A. The side chain of K4089 forms a salt bridge with the side chain of D205 and establishes a hydrogen bond with the oxygen atom in the main chain of N203. Q222 and D205 of Lis1 also form polar interactions with S4115 and Q4117 and N4085 residues of dynein (**Fig. 4e, Extended Data Fig. 8b**).

The AAA5-Lis1_ring_ WD-40 interface shows a more compact interaction (**Fig. 4f**). This interface is mainly composed of residues Q3636, S3613, T3612, D3616, D3617, A3618, and K3621 of dynein MD-A and Y225, C226, G190, H191, M172, and D192 of the Lis1 WD-40 domain (**Fig. 4f and Extended Data Fig. 8b**). Notably, the side chain of Y225 of Lis1 forms a polar interaction with the side chain of Q3636. Additionally, the side chain of D3616 and the main chain of D3617 form hydrogen bonds with the main chain of C226. The side chain of A3618 forms hydrophobic interaction with the main chain of G190, while residues M172 and H191 of Lis1 form hydrophobic and hydrophilic interactions with the side chain of K3621. Additionally, novel interactions are formed between MD-A and MD-B in the Phi^L^-Lis1, compared with canonical Phi (**Extended Data Fig. 10**), suggesting that the dynein Phi^L^-Lis1 is a stable conformation.

### The Phi^L^-Lis1 interface regulates Nde1-dependent dynein activation

To evaluate whether the new interaction sites we detected between Lis1 and dynein MD-A in Phi^L^-Lis1 are critical for activation of dynein, we introduced three sets of Lis1 mutations targeting the interfaces that interact with the linker, AAA6, and AAA5 of dynein. Key residues of Lis1_ring_ at each interface were mutated to charged residues or alanine to disrupt these interactions (Lis1^linker^: E300K, K303E, S304R and M329A; Lis1^AAA6^: N203K, D205K and Q222A; Lis1^AAA5^: M172K, D192K, Y225A, C226D). Similar to wild-type Lis1 (WT Lis1), these Lis1 mutants formed homodimers and interacted with Nde1 (**Fig. 4g**). However, the binding efficiency of Nde1 was reduced for the Lis1^AAA5^ mutant (**Fig. 4g**), suggesting that the AAA5-Lis1_ring_ interface may share a region involved in Nde1 binding to Lis1 (**Extended Data Fig. 11**). The Lis1 mutants also bound to dynein, and the dynein binding efficiency of Lis1 was increased with Nde1 (**Fig. 4h**). Notably, we detected a mass population corresponding to dynein bound to one Nde1 and two Lis1^AAA6^ mutants, indicating that mutations to the AAA6 interaction site of Lis1 prevent dissociation of Nde1 from the dynein-Lis1 complex (**Fig. 4h**).

To determine how these mutations affect activation of dynein motility, we assayed single molecule motility of complexes assembled with wild type dynein, dynactin, and the BicDR1 adaptor (DDR) on surface-immobilized MTs in vitro in the presence and absence of Lis1 and Nde1. Consistent with our previous observations^26^, Lis1 enhanced the run frequency of DDR about 3-fold, and Nde1 and Lis1 together increased the run frequency 15-fold (**Fig. 4i, j and Supplementary Video 2**). In the absence of Nde1, Lis1 mutants activated dynein motility at similar levels of WT Lis1. In the presence of Nde1, Lis1^linker^ and Lis1^AAA5^ triggered activation of DDR motility similar to WT Lis1 (**Fig. 4i, j**). In comparison, Nde1 failed to enhance Lis1^AAA6^-mediated dynein motility, suggesting that Lis1^AAA6^ cannot form the stable Phi^L^-Lis1 complex and open the Phi conformation (**Fig. 4i, j**). Together with MP results, we demonstrate that mutations in the Lis1^AAA6^ interface disrupt Nde1-mediated opening of the Phi conformation by Lis1, highlighting the importance of the Phi^L^-Lis1 structure in the dynein activation pathway.

## Discussion

In this study, we investigated the structure and mechanism of how Lis1 and Nde1 rescue dynein from autoinhibition prior to the assembly of active dynein transport machinery. Using negative stain EM imaging, we directly showed that Nde1 and Lis1 cooperatively promote the opening of Phi dynein, whereas neither Nde1 nor Lis1 alone exhibited a significant effect on Phi opening (**Fig. 1c-e**). Lis1 can readily bind to open dynein (**Extended Data Fig. 1**) and facilitate the assembly of DDA complexes and Nde1 addition does not further enhance DDA motility^26^. Despite that Lis1 alone can bind to the Phi dynein and potentially open this autoinhibited conformation^45^, we show that Nde1 facilitates Lis1 to dock onto Phi dynein more efficiently and promote its switch it to the open state before DDA assembly in this work. Similar to molecular chaperones, Nde1 rapidly dissociates from dynein after handing off Lis1, promoting dynein-Lis1 complex formation but not existing in the final complex. Because Nde1 has an overlapping binding site with the p150 subunit of dynactin on DIC-N^12^, dissociation of Nde1 from DIC-N after tethering Lis1 to dynein may enable efficient recruitment of dynactin to dynein-Lis1 complexes.

Our structural and functional studies of dynein-Lis1 complexes revealed a key intermediate step on Lis1 and Nde1 mediated opening of dynein. Although a dynein dimer contains Lis1 binding sites on each motor domain, our MP assays showed that Phi dynein recruits a single Lis1. Using cryo-EM imaging, we revealed a Phi^L^-Lis1 structure in which the two AAA+ rings of Phi dynein rotate slightly backward to accommodate Lis1 binding to the front side. This rotational motion reduces the spacing between the AAA+ rings, thereby preventing Lis1 binding to the back side. Preferential binding of Lis1 to the front side may be due to the IC-LC tower, which is located at the front side of the Phi motor.

The Phi^L^-Lis1 is fundamentally distinct from the previously reported Chi of yeast dynein monomers^37^. The Chi is stabilized by two Lis1, one on each side, and adopts a more open and extended conformation compared to the Phi^L^-Lis1 motor domains (**Extended Data Fig. 12**). In comparison, our study utilized full-length, wild-type human dynein and we could not detect Chi dynein even when we used this conformation as a reference during cryo-EM image processing. It is possible that isolated dynein motor domains may prefer to recruit two Lis1s and form more extended Chi. In comparison, full-length dynein readily forms the compact Phi and structural constraints imposed by the tail domain may restrict the relative movement of the Phi motor domains. Lis1 binding to the front side of Phi^L^ dynein reduces the spacing on the back side, thereby preventing the formation of Chi. Most interaction sites located at Lis1_ring_ surface of Chi-Lis1 are also present in that of Phi^L^-Lis1 (**Extended Data Fig. 9**). In comparison, Phi^L^-Lis1 exhibits more compact interactions (**Fig 4 and Extended Data Fig. 8,10**). Consistent with Phi^L^-Lis1, we did not detect complexes with one dynein and two Lis1s in MP, suggesting that Phi^L^-Lis1, rather than Chi, is the stable intermediate of full-length human dynein.

The mutagenesis of the interactions between Lis1 and the AAA6 subdomain of MD-A in Phi^L^-Lis1 disrupts Nde1’s ability to promote Lis1-mediated dynein activation, confirming that the Phi^L^-Lis1 is a key intermediate in the dynein activation pathway. However, these mutations did not disrupt the mutant Lis1’s ability to increase the run frequency of dynein several-fold on its own. This is because WT dynein can be either in open or Phi conformations with near equal probability in our conditions. Our model predicts that mutant Lis1 can still bind and enhance DDA assembly of open dynein without Nde1. However, it cannot further enhance dynein motility synergistically with Nde1 because this mutant is deficient in forming Phi^L^-Lis1.

Based on our results and previous observations, we propose a mechanism underlying dynein activation by Lis1 and Nde1. In the absence of Nde1, Lis1 alone can bind to both DIC-N and dynein motor domains^11^, inducing a conformational change from canonical Phi to Phi^L^. However, the efficiency of this process is low and Lis1 cannot open Phi^L^ dynein on its own (**Fig.1c-e, 2a**). In the presence of Nde1, Lis1-Nde1 is initially recruited to DIC-N positioned at the front side of Phi dynein, facilitating more efficient binding of Lis1 to the front side of Phi. The local enhancement of Lis1 near the Phi motor by Nde1 facilitates more efficient binding of Lis1 to Phi dynein (**Fig. 5 step-i, -ii**). Nde1 and Lis1 form a transient Phi^L^-Lis1-Nde1 complex (**Fig. 5 step-i**). Nde1 dissociates spontaneously, leading to Phi^L^-Lis1 formation (**Fig. 5 step-ii**). Lis1 binding induces a slight backward rotation of the two motor rings in Phi^L^-Lis1 (**Fig. 3c**), suggesting an intermediate state prior to an open state. Additionally, a slight twist in the neck region, caused by motors rotation and likely inducing an unwinding trend in the tail, may also contribute to dynein opening (**Fig. 3d**). Subsequently, Phi^L^-Lis1 transitions to open dynein-Lis1 with the assistance of Nde1 (**Fig. 5 step-iii**). The binding of Lis1 to dynein facilitates DDA assembly and activates dynein motility by recruiting the p150 subunit of dynactin to dynein through its LisH domain^11^ (**Fig. 5, step-iv**). Future studies are required to understand how Nde1 hands off Lis1 to dynein and why it rapidly dissociates from dynein. In addition to its tethering role and facilitating the formation of Phi^L^-Lis1, it remains to be determined whether Nde1 has additional roles in helping Lis1 convert Phi^L^-Lis1 to the open dynein.

**Fig. 5.**
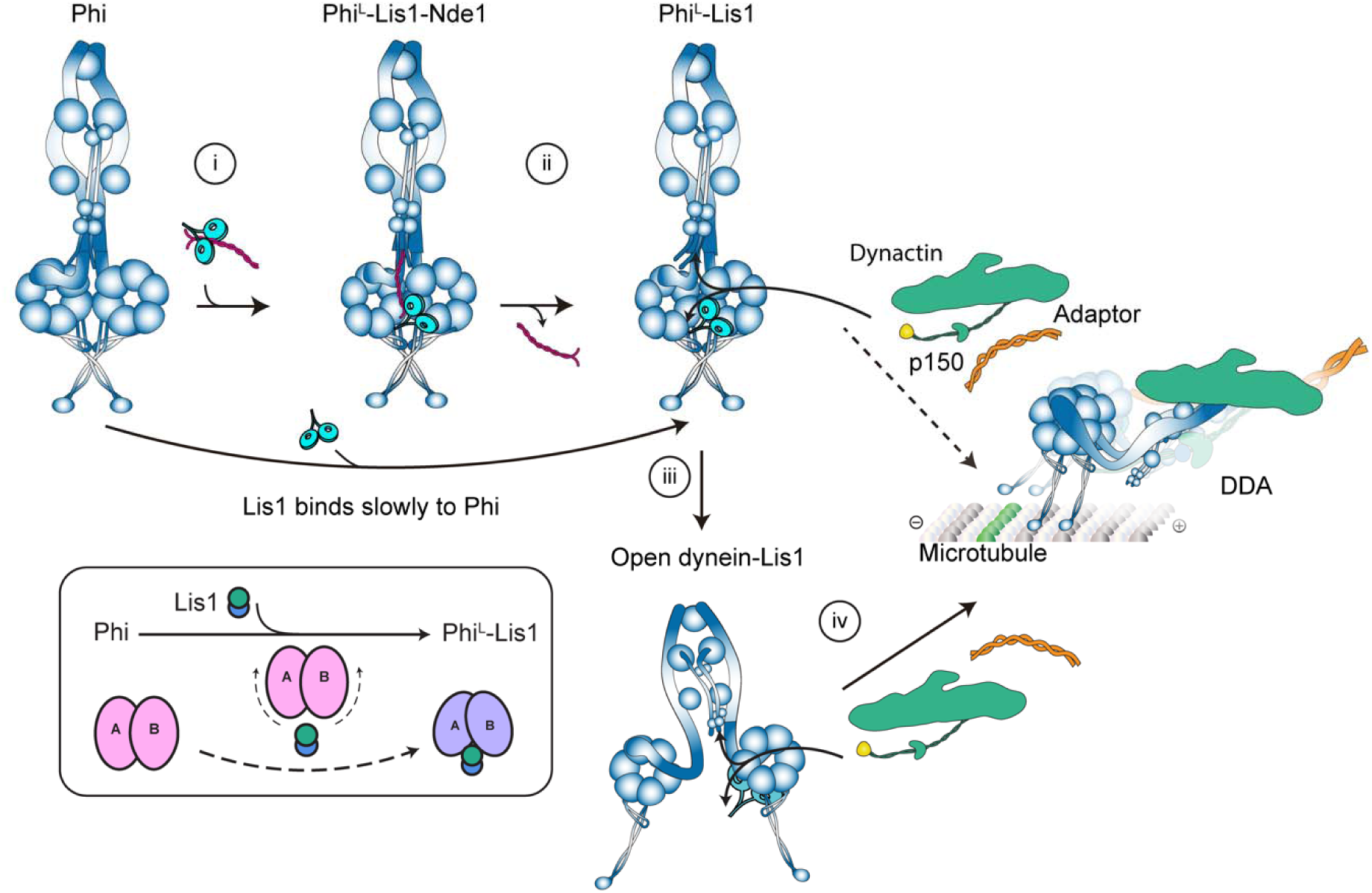
Model for the initial stage of dynein activation by Nde1 and Lis1. **Step-i,** Nde1 tethers Lis1 to Phi dynein and forms a transient Phi^L^-Lis1-Nde1 complex. **Step-ii,** Nde1 dissociates spontaneously from dynein, leading to the formation of Phi^L^-Lis1. Without Nde1, Lis1 binding to Phi dynein and formation of Phi^L^-Lis1 becomes a less efficient and slow process. The binding of Lis1 induces a slight backward rotation of the two motor rings and a twist in the neck region, likely leading an unwinding trend in the tail and contributing to dynein opening. **Step-iii**, Phi^L^-Lis1 transitions to open dynein-Lis1 assisted by Nde1. **Step-iv**, Lis1 promotes processive DDA complex assembly by interacting with p150 and DIC-N through its LisH domain. The dashed line indicates the possibility that Lis1 in the Phi^L^-Lis1 complex promotes dynactin recruitment and DDA assembly without switching to the open conformation.

## Methods

### Cloning and expression

The plasmid encoding full-length human dynein^63^ was generously provided by Andrew Carter (His-ZZ-TEV-SNAPf DHC1_IC2C_LIC2_Tctex1_Robl1_LC8, Addgene plasmid #111903). The His-ZZ-TEV-SNAPf tag is fused to the N-terminus of the dynein heavy chain. Human Lis1 and Nde1 (residues N-terminal 1-190 residues, which functions similarly to full-length Nde1 in the single motility assay^26^), and the mouse BIDCR1 gene, were each cloned individually into the pOmniBac backbone. The constructs featured a ZZ-TEV tag at the N-terminus and a SNAPf tag at the C-terminus. Lis1 mutants containing point mutations were generated using purchased DNA fragments (IDT) containing the mutations and inserted into the plasmid backbone. The mutations were verified by Oxford Nanopore full-plasmid sequencing. The constructs used in this study are listed in **Supplementary Table 1**.

These proteins were all expressed in insect sf9 cells, as describe previously^14,19,26^ with slight modifications. Briefly, Bacmid DNA isolated from the from DH10MultiBac competent cells (Geneva Biotech) were transfected into the in sf9 insect cells with the Cellfectin® II (Gibco) reagent. Protein expression in sf9 cells was accomplished by infecting them with P2 virus at a cell density of 2.5 million cells/mL. For dynein expression, 28 mL of P2 virus was added into a 1.4 L culture of sf9 cells. For Lis1, Nde1, and BicDR1, 7 mL P2 virus was used to infect the 0.7 L sf9 cells. Cells were harvested after 75 hours by centrifugation at 1000 rcf for 15 minutes at 4°C. The cell pellets were flash-frozen in liquid nitrogen and stored at -80°C.

### Protein purification

Purification for full-length human dynein was previous described^19^. Briefly, the cell pellets from a 1.4 L cell culture were resuspended in 100 mL lysis buffer (50 mM HEPES pH 7.2, 100 mM NaCl, 1 mM DTT, 0.1 mM ATP, 10% glycerol) containing 2 tablets of Complete EDTA-free protease inhibitor (Roche) and 2 mM PMSF. The suspension was homogenized using a Dounce with a tight plunger for 15∼25 strokes, followed by clarification through centrifugation at 65,000 rpm with a Ti70 rotor (Beckman) for 1 hour at 4°C. The supernatant was then incubated with 3 mL lgG Sepharose 6 fast flow resin (Cytiva) for 3∼4 hours on a roller at 4°C, followed by washed with 200 mL lysis buffer and 200 mL TEV buffer (50 mM Tris–HCl pH 7.4, 150 mM K-acetate, 2 mM Mg-acetate, 1 mM EGTA, 10% glycerol, 0.1 mM ATP, 1 mM DTT). Afterward, the resins were incubated with TEV buffer supplemented with 400 ug TEV protease overnight at 4°C. The supernatant was collected and concentrated with a 100 kDa MWCO Amicon concentrator, then loaded into a TSKgel G4000 column pre-equilibrated with the GF150 buffer (25 mM HEPES pH 7.2, 150 mM KCl, 1 mM MgCl_2_, 5 mM DTT, 0.1 mM ATP). Peak fractions were collected and concentrated to 2∼3 mg/mL for Cryo-EM grid preparation. The quality of the sample was evaluated with the SDS-PAGE gels and the negative-stain EM.

The purification of Lis1, Nde1, and BicDR1 from a 0.7 L cell culture followed a similar protocol to that of dynein, with a few modifications. Specifically, 50 mL of lysis buffer was used to resuspend the cell pellets, and ATP was omitted from the GF150 buffer. And Superose 6 Column (Cytiva) was used for size exclusion chromatography. The concentrated proteins were aliquoted, flash-frozen in liquid nitrogen, and stored at -80°C. The quality of the proteins was assessed using SDS-PAGE gels.

Dynactin was isolated from pig brains through a series of purification steps, including SP Sepharose Fast Flow and MonoQ ion exchange chromatography (Cytiva), followed by size exclusion chromatography using a TSKgel G4000SWXL column (Tosoh), as described by previous protocol^64^.

### MT reconstitution

MTs were reconstituted using porcine tubulin, which was either purchased from Cytoskeleton or purified in-house in MT buffer (25 mM MES, 70 mM NaCl, 1 mM MgCl2, 1 mM EGTA, and 1 mM DTT, pH 6.5). The tubulin was concentrated to 10 mg/mL at 4°C, then flash-frozen and stored at -80°C. To polymerize the MTs, the tubulin was diluted to 5 mg/mL in MT buffer supplemented with 3 mM GTP. The tubulin mixture was incubated on ice for 5 minutes and then transferred to a 37°C incubator for 1 hour. After incubation, the MTs were pelleted at 20,000 rcf for 8 minutes at room temperature and resuspended in MT buffer supplemented with 5 μM paclitaxel before being stored at room temperature.

### MP assay

High-precision coverslips (Azer Scientific) were cleaned by alternating washes with isopropanol and water three times in a bath sonicator, then air-dried. The gasket was cleaned similarly, without sonication, and air-dried before being placed onto a clean coverslip. A total of 14 µL of filtered mass photometry buffer (30 mM HEPES pH 7.4, 5 mM MgSO_4_, 1 mM EGTA, and 10% glycerol) was added to a well for autofocus. The protein sample was then applied to the well and diluted to a concentration of 5–20 nM in the buffer. Protein contrast data were collected using a TwoMP mass photometer (Refeyn 2) with two technical replicates. The instrument was calibrated with a standard mix of conalbumin, aldolase, and thyroglobulin. MP profiles were analyzed by fitting to multiple Gaussian peaks, with the mean, standard deviation, and percentages calculated using DiscoverMP software (Refeyn). The data for parameters of a multi-Gaussian fit of MP measurements is summarized in **Supplementary Table 2**.

### Negative-stain EM and data quantification

Freshly purified dynein was diluted in GF150 buffer to a final concentration of 14.3 nM in the presence of 0.1 mM ATP and subsequently evaluated using negative-stain electron microscopy (EM). A 4 µL aliquot of the sample was applied to glow-discharged carbon film grids (Electron Microscopy Sciences) and stained with 2% uranyl acetate. The grids were then imaged using a 120 kV Talos L120C electron microscope. Micrographs were manually acquired at a magnification of 45,000x. More than 40 micrographs were collected per experiment for each condition to obtain sufficient particles for statistical analysis.

Samples for statistical analysis of the Phi ratio were prepared with the following molar ratios: dynein: Lis1(dimer) at 1:0, 1:1, and 1:2; dynein: Nde1(dimer) at 1:0, 1:1 and 1:2; dynein : Lis1 : Nde1 at 1:0:0, 1:1:1 and 1:2:2. The mixtures were incubated on ice for 90 minutes and subsequently subjected to negative staining. Each experimental group was accompanied by its respective control. The assay was repeated independently three or more times, using different batches of freshly purified protein.

Micrographs were processed using cryoSPARC, including blob picking, micrograph extraction, and 2D classification. Briefly, micrographs from each experimental group were merged, and particle picking was performed using templates of Phi and open dynein. The particle diameter was set to 750 Å, and the distance cutoff for dynein particles was 400 Å to optimize particle selection. All particles were extracted, followed by three rounds of 2D classification. Phi and open dynein particles were identified from the 2D classification and traced back to the corresponding micrographs for each condition, where Phi and open particles were quantified. To calculate the normalized fraction of Phi, the total number of Phi and paired open dynein particles was determined for micrographs under a given condition. The ratio of Phi particles in the dynein-alone condition (N(Phi) / N(Phi + open)) was defined as proportion A, serving as the control for each group. The ratio of Phi particles in each experimental condition, excluding the dynein-alone control, was defined as proportion B (N(Phi) / N(Phi + open)). The normalized fraction of Phi was calculated as B/A, and GraphPad Prism was used to plot the normalized fraction of Phi.

### Cryo-EM sample preparation

For the dynein, Lis1, and Nde1 sample, Lis1 and Nde1 were incubated at a 1:1 molar ratio for 30 minutes on ice. Freshly purified dynein, at a concentration of 2 mg/mL, was then added to the Lis1-Nde1 complex at a 1:2:2 molar ratio and incubated on ice for 5 minutes, with 5 mM ATP added immediately prior to freezing. For the dynein and Lis1 complex, dynein (2 mg/mL) was incubated with Lis1 for 90 minutes on ice, and 5 mM ATP was added just before vitrification.

For vitrification, 3.5 μL of the prepared sample was applied to glow-discharged Quantifoil holey carbon grids (R2/1, 300 mesh gold), which were treated for 45 seconds at 25 mA using a GloQube Glow Discharge system (Quorum Technologies). The grids were blotted for 2.5 to 4.5 seconds at 4°C and 100% humidity, then vitrified by plunging into liquid ethane using a Vitrobot Mark IV (Thermo Fisher Scientific).

### Cryo-EM data collection

Data were collected at the Yale ScienceHill Cryo-EM facility using a Glacios microscope (Thermo Fisher Scientific) operated at 300 keV and equipped with a K3 detector. Data collection was facilitated by SerialEM software, targeting a defocus range of -1.2 μm to - 2.6 μm. Four exposures per hole were recorded as movies, comprising 40 frames each, with a total electron dose of 40 e⁻/Å². A total of 7,128 movies were collected for the Dynein-Lis1-Nde1 condition, while 16,558 movies were collected for the Dynein-Lis1 condition.

### Cryo-EM data processing

Cryo-EM movies were pre-processed using CryoSPARC Live, which included patch motion correction and patch CTF estimation. The processing workflows are illustrated in **Extended Data Fig. 4 and 5**. The statistics are summarized in **Table 1**.

**Table 1.**
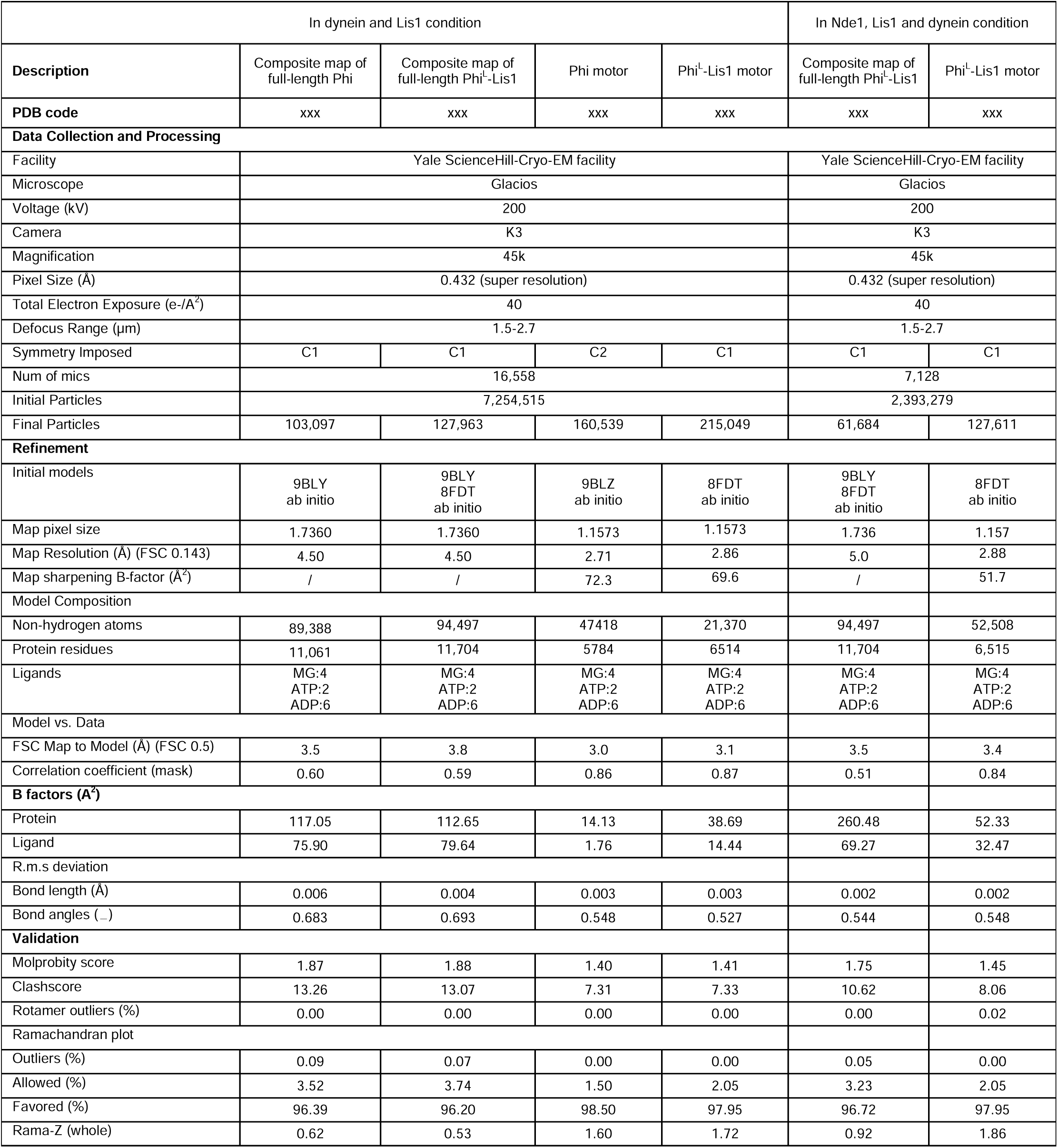
Cryo-EM data collection, refinement, and validation statistics.

For the dynein-Lis1 condition dataset, particles were picked using the blob picker, extracted with a box size of 512 pixels, and downscaled to 128 pixels with a pixel size of 3.456 Å. In total, 7,254,515 particles were extracted. The iterative 2D classification was performed to filter the particles, resulting in the selection of 204,005 high-quality particles for ab initio reconstruction. Initial maps for the dynein single motor domain and the Phi^L^-Lis1 motor domains were identified. The map of the Phi^L^-Lis1 motor domains was subsequently used for heterogeneous refinement of all original particles. The original particles were divided into four subsets, each subjected to heterogeneous refinement (4 classes). Three rounds of heterogeneous refinement were performed, updating the reference each time, ultimately identifying the Phi and Phi^L^-Lis1 motor domains. The Phi and Phi^L^-Lis1 motor domains were merged separately and extracted from micrographs using a box size of 512 pixels, which was then binned to 384 pixels, resulting in a pixel size of 1.1573 Å. Two rounds of heterogeneous refinement were conducted to exclude junk particles. High-quality subsets were selected for homogeneous refinement, followed by two rounds of CTF refinement and local refinement. The Phi^L^-Lis1 motor domains achieved a resolution of 2.86 Å, and the Phi motor domains reached a resolution of 2.71 Å, exhibiting C2 symmetry.

To reconstitute the tails of the Phi^L^-Lis1 and Phi motor domains, the particles were recentered at the tail and then extracted from micrographs using a box size of 512 pixels, binned to 256 pixels, yielding a pixel size of 1.736 Å/px. Following this, heterogeneous refinement was applied to filter the particles, and high-quality subsets were selected for homogeneous refinement. The overall tail resolutions reached 4.21 Å for the Phi tail and 4.05 Å for the Phi^L^-Lis1 tail. Four masks were devised to cover the tail segments, which were divided into the NDD, left, right, and neck regions. Maps of local refinement using these masks were integrated, referring to the consensus map of the tail. The full-length map was assembled by stitching together the tail and motor domains in ChimeraX, corresponding to Phi and Phi^L^-Lis1, respectively.

For the Lis1-Nde1-dynein condition dataset, the process closely mirrored that of the dynein-Lis1 condition described above. Briefly, the blob picker identified 2,393,279 particles. Following extraction from the micrographs and iterative 2D classification, 204,005 particles were selected for initial map generation through ab initio reconstruction. An initial map for the Phi^L^-Lis1 motor domains were obtained, which were then subjected to iterative heterogeneous refinement using all particles with the Phi^L^-Lis1 initial map as a reference. However, the Phi motor domains did not appear in the heterogeneous refinement, even when the verified Phi map from this study was utilized as a reference. Subsequently, the Phi^L^-Lis1 domain map was re-extracted from the micrographs and underwent two rounds each of heterogeneous refinement, CTF refinement, and local refinement, achieving a resolution of 2.88 Å. The consensus map of the tail reached a resolution of 6.22 Å after recentering and re-extracting the tail region. Masks and local refinement were employed to enhance the local resolution of the tail. The full-length Phi^L^-Lis1 structure was reconstituted by integrating the composite tail map and motor map in ChimeraX.

### Model building and refinement

For model building, previously reported structures 9BLY^19^, 9BLZ^19^, and 8FDT^38^ were utilized as the initial models for the full-length and motor domains of Phi and Phi^L^-Lis1. The individual domains, including the tail, single motor and Lis1 dimer, were extracted from the 9BLY, 9BLZ, and 8FDT, and rigid-body fitting into the Cryo-EM maps were performed using UCSF ChimeraX. The models were then manually constructed in COOT^65,66^, and followed by real-space refinement in Phenix^67^. The quality of the refined models was assessed using the MolProbity integrated into Phenix, with the statistics reported in **Table 1**.

### Single-molecular motility assay

Fluorescent imaging was conducted using a custom-built, multicolor objective-type TIRF microscope based on a Nikon Ti-E microscope body. It was equipped with a 100X magnification, 1.49 N.A. apochromatic oil-immersion objective (Nikon) and a Perfect Focus System. Fluorescence signals were captured by an electron-multiplied charge-coupled device camera (Andor, Ixon EM+, 512 × 512 pixels), with an effective pixel size of 160 nm after magnification. Probes such as Alexa488/GFP/mNeonGreen, LD555, and LD655 were excited by 488 nm, 561 nm, and 633 nm laser beams (Coherent), coupled to a single-mode fiber (Oz Optics), and their emissions were filtered using 525/40, 585/40, and 697/75 bandpass filters (Semrock), respectively. The entire system was controlled via MicroManager 1.4 software.

Biotin-PEG treated flow chambers were treated with 5 mg/ml streptavidin for 2 minutes, followed by washing with MB buffer (30 mM HEPES pH 7.0, 5 mM MgSO₄, 1 mM EGTA, 1 mg/ml casein, 0.5% pluronic acid, 0.5 mM DTT, and 1 µM Taxol). Biotinylated MTs were then added to the chamber for 2 minutes and washed again with MB buffer. Proteins were prepared by diluting them to the desired concentrations in MB buffer. For DDRNL complex assembly, a mixture of 10 nM dynein, 150 nM dynactin, 50 nM BicDR1, 200nM Lis1 and 10nM Nde1 was incubated on ice for 15 minutes, then diluted tenfold into imaging buffer (MB buffer containing 0.1 mg/ml glucose oxidase, 0.02 mg/ml catalase, 0.8% D-glucose, and 2 mM ATP) and introduced to the flow chamber. Motility was observed and recorded for 40 seconds.

### Data analysis for single-molecular motility assay

Single-molecule motility of the DDR complex was captured for 200 frames per imaging area and analyzed as kymographs made in FIJI. Run frequency was determined by counting the number of processive BicDR1 molecules on each MT, then dividing this number by the MT length and the total data collection time, with a custom MATLAB script. The p-values for the two-tailed Student’s t-test were determined in Excel.

## Data availability

Cryo-EM Density maps and models have been deposited in the Electron Microscopy Data Bank and Protein Data Bank as follows: In the dynein and Lis1 condition: PDB-XXX/EMD-XXX for full length Phi and PDB-XXX/EMD-XXX for motor domains of Phi; PDB-XXX/EMD-XXX for full length Phi^L^-Lis1 and PDB-XXX/EMD-XXX for motor domains of Phi^L^-Lis1. In the dynein, Lis1 and Nde1 condition: PDB-XXX/EMD-XXX for full-length Phi^L^-Lis1 and XXX/EMD-XXX for motor domains of Phi^L^-Lis1.

## Acknowledgement

We are very grateful to members of the Zhang and Yildiz laboratories for their valuable discussions. This work was funded by the NIH/NIGMS (GM136414 to A.Y., and GM142959 to K.Z.) and in part by a Collaboration Development Award Program (to K.Z.) from the Pittsburgh Center for HIV Protein Interactions (U54AI170791). The Cryo-EM data were collected at the Yale ScienceHill Cryo-EM facility. We would like to thank Jianfeng Lin and Kaifeng Zhou for assistance with the data collection. Yale Cryo-EM Resource is funded in part by the NIH grant S10OD023603 awarded to Frederick J. Sigworth.

## Author contributions

K.Z. and A.Z. designed the study. J.Y. expressed and purified dynein, Lis1 and Nde1 proteins for EM. J.Y. and P.C. prepared the cryo-EM samples, collected and processed the data, and built the PDB models. P.C. and J.Y. processed negative stain EM data and quantified the particle numbers. Y.Z. performed Lis1 mutagenesis, protein preparation, TIRF and mass photometry assays. J.Y., P.C., Y.Z., K.Z., and A.Y. analyzed the data and prepared figures. J.Y., Y.Z., P.C., K.Z., and A.Y. wrote the manuscript with input from all authors.

## Competing Interests

The authors declare no competing interests.

## Figures

**Supplementary Table 1.**
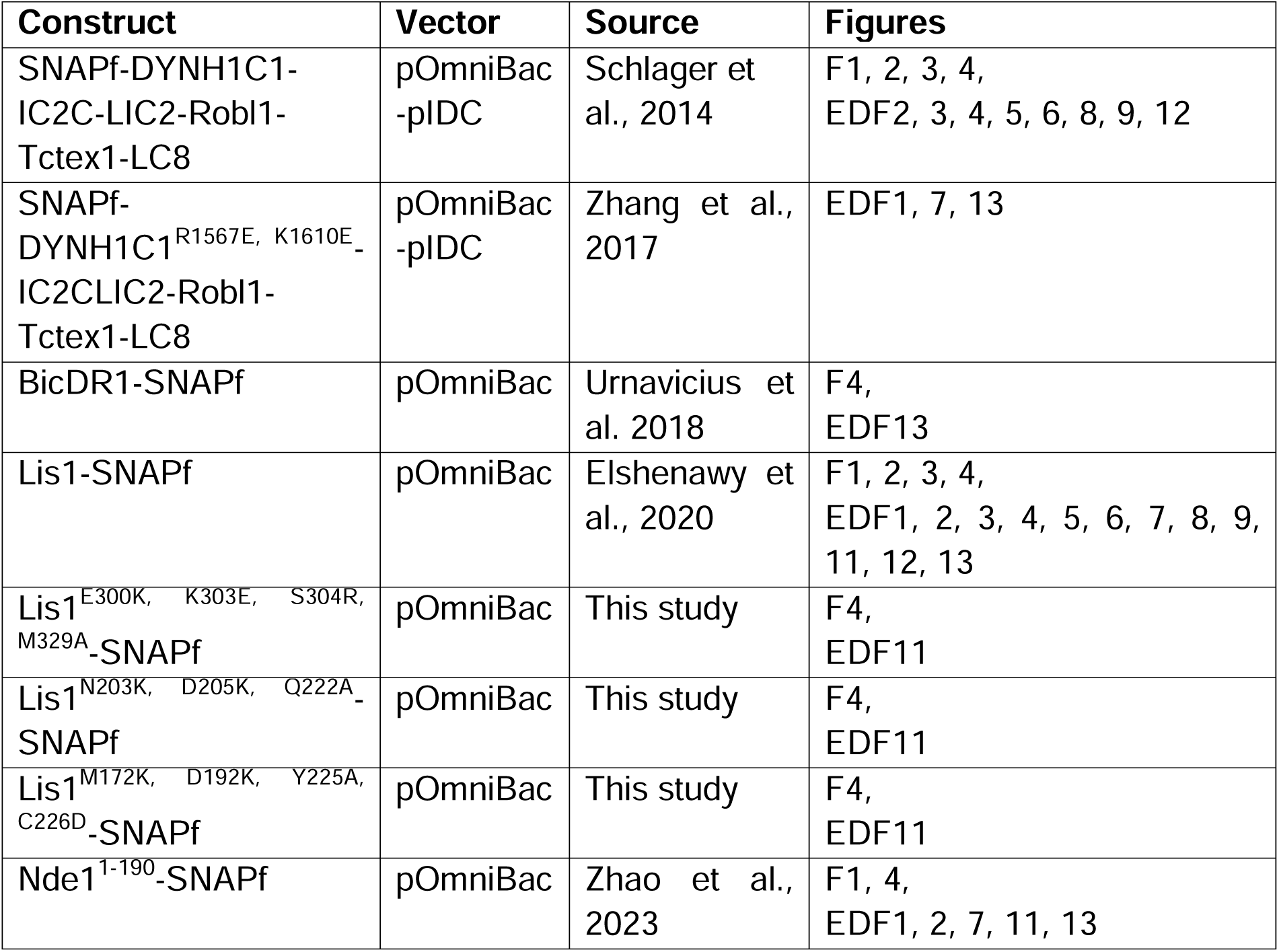
The list of protein constructs used in this study. Dynein chains were codon-optimized for *Spodoptera frugiperda* (Sf9) expression and inserted into the pOmniBac backbone. Nde1, Lis1, and BicDR1 constructs were cloned into the pOmniBac backbone. Constructs were tagged with an N-terminal 6xHis-ZZ-TEV site for affinity purification and TEV protease cleavage during protein purification. The SNAPf tag was inserted for labeling the proteins with fluorescent dyes (F: Figure, EDF: Extended Data Figure).

| Construct | Vector | Source | Figures |
| --- | --- | --- | --- |
| SNAPf-DYNH1C1-IC2C-LIC2-Robl1-Tctex1-LC8 | pOmniBac-pIDC | Schlager et al., 2014 | F1, 2, 3, 4, EDF2, 3, 4, 5, 6, 8, 9, 12 |
| SNAPf-DYNH1C1<sup>R1567E, K1610E</sup>-IC2CLIC2-Robl1-Tctex1-LC8 | pOmniBac-pIDC | Zhang et al., 2017 | EDF1, 7, 13 |
| BicDR1-SNAPf | pOmniBac | Urnavicius et al. 2018 | F4, EDF13 |
| Lis1-SNAPf | pOmniBac | Elshenawy et al., 2020 | F1, 2, 3, 4, EDF1, 2, 3, 4, 5, 6, 7, 8, 9, 11, 12, 13 |
| Lis1<sup>E300K, K303E, S304R, M329A</sup>-SNAPf | pOmniBac | This study | F4, EDF11 |
| Lis1<sup>N203K, D205K, Q222A</sup>-SNAPf | pOmniBac | This study | F4, EDF11 |
| Lis1<sup>M172K, D192K, Y225A, C226D</sup>-SNAPf | pOmniBac | This study | F4, EDF11 |
| Nde1<sup>1-190</sup>-SNAPf | pOmniBac | Zhao et al., 2023 | F1, 4, EDF1, 2, 7, 11, 13 |

**Supplementary Table 2.**
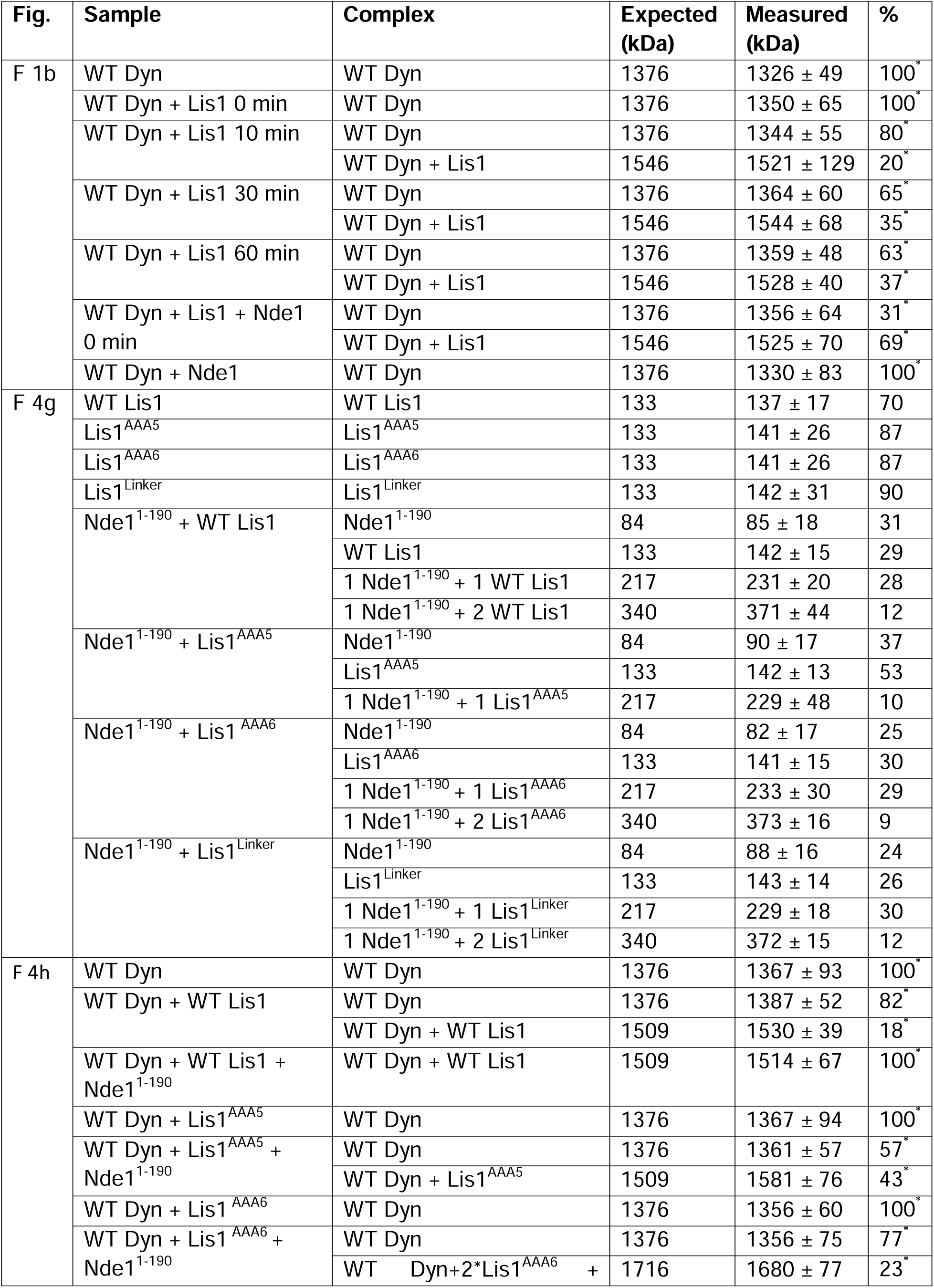
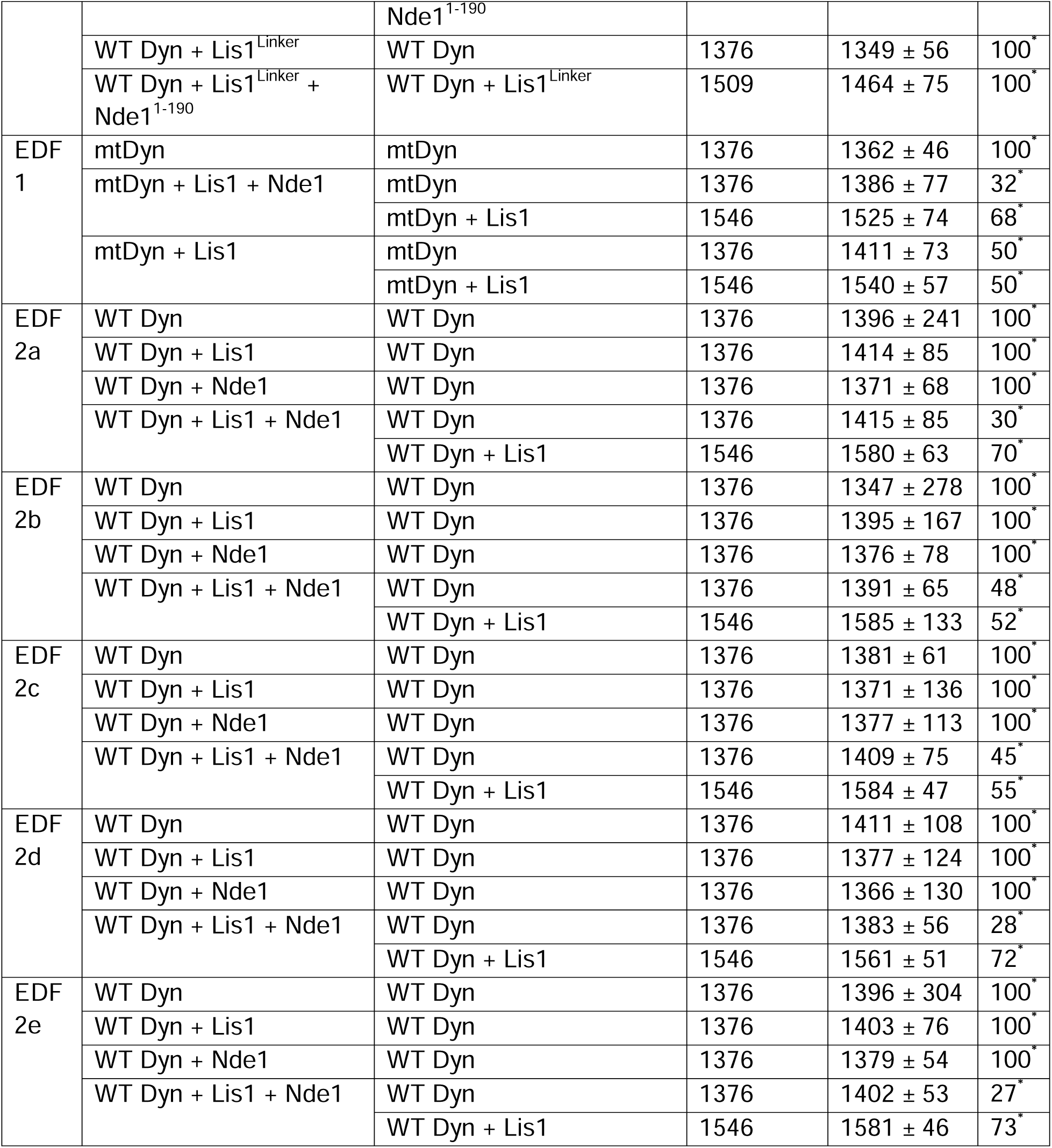
The parameters of a multi-Gaussian fit of MP measurements. Dynein (Dyn), Lis1, and Nde1 were mixed at 1:2:2 ratio if present and diluted into 5-20 nM. Measured mass and percentage represent the center (mean ± s.d.) and the percent area of the corresponding Gaussian peak (*percentages are normalized so that the sum of all dynein included peaks are 100%). Expected mass corresponds to the dimeric forms of Lis1, Nde1, and Dyn (F: Figure, EDF: Extended Data Figure). Lis1 used in F 1b, and EDF 1 contains a ZZ tag, and its expected MW is 170 kDa.

## Supplementary Video Legends

**Supplementary Video 1.** Full-length human dynein in Phi^L^ conformation, bound to a Lis1 dimer and displaying the newly identified interface with Lis1.

**Supplementary Video 2.** Single-molecule motility recordings of WT DDR complexes, in the presence or absence of Nde1, WT Lis1, and Lis1 mutants. The fluorescence signal originates from BicDR1-mNeonGreen.

## Extended Data Figures

**Extended Data Fig. 1.**
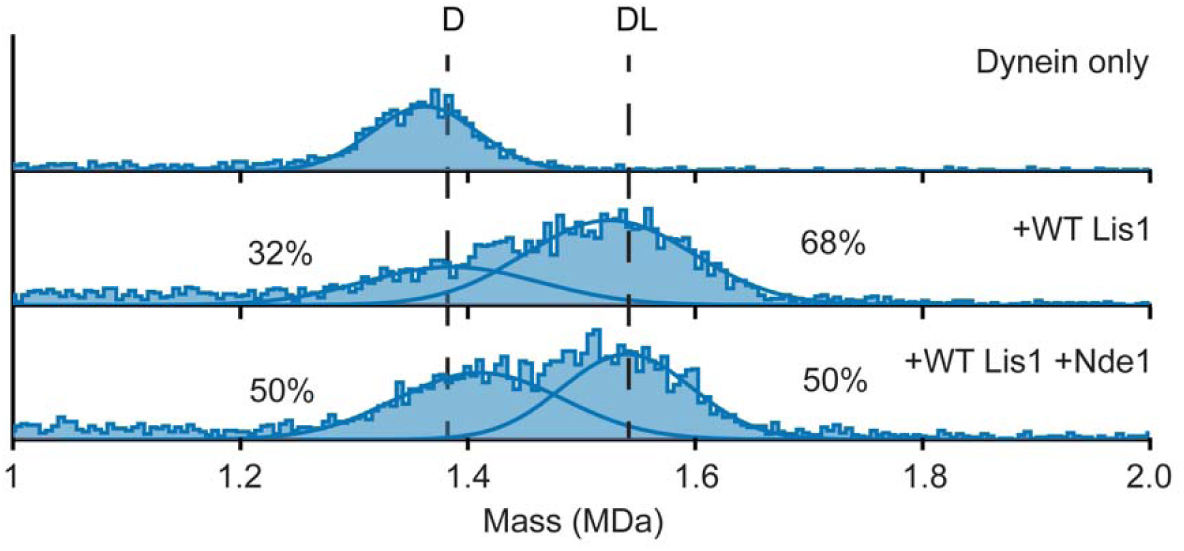
MP analysis of Nde1’s effect on Lis1 binding to open dynein. MP shows that Nde1 does not promote increased Lis1 binding to open dynein compared to the Lis1-alone condition, indicating that Lis1 can efficiently bind to open dynein and Nde1 does not enhance Lis1’s interaction with open dynein.

**Extended Data Fig. 2.**
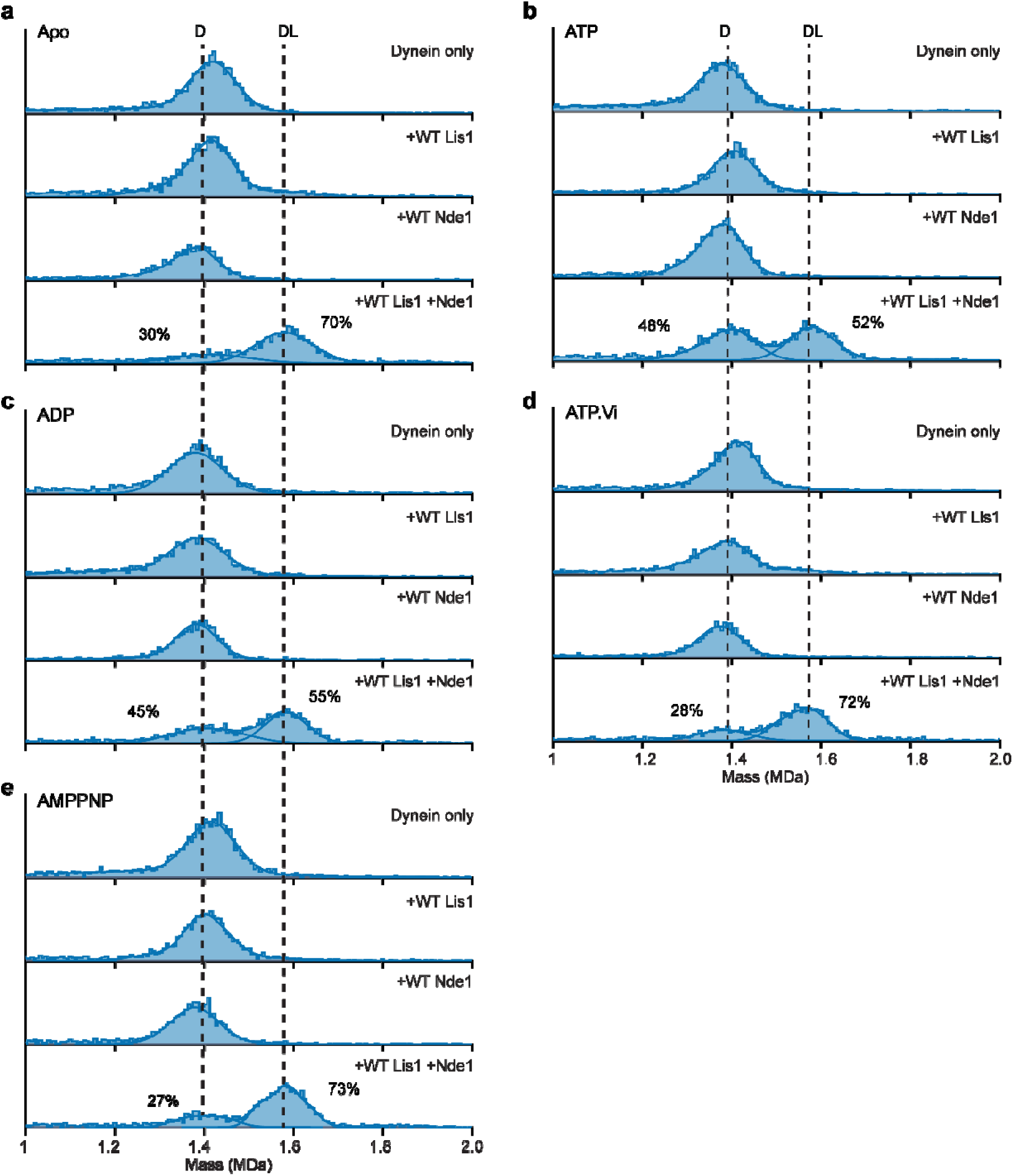
MP analysis of nucleotide conditions on Nde1 and Lis1 binding to WT dynein. MP shows that under apo buffer (**a**), 0.1 mM ATP (**b**), ADP (**c**), ATP.vi (**d**), and AMPPNP (**e**) conditions, Nde1 promotes Lis1 binding to dynein, forming a 1:1 dynein-Lis1 (DL) complex. The nucleotide condition does not affect Nde1’s ability to tether Lis1 to dynein. Importantly, the formation of dynein-Nde1, dynein-Lis1-Nde1 and 1:2 dynein-Lis1 complexes were not observed. In the Lis1-alone condition, no significant DL complex was formed immediately.

**Extended Data Fig. 3.**
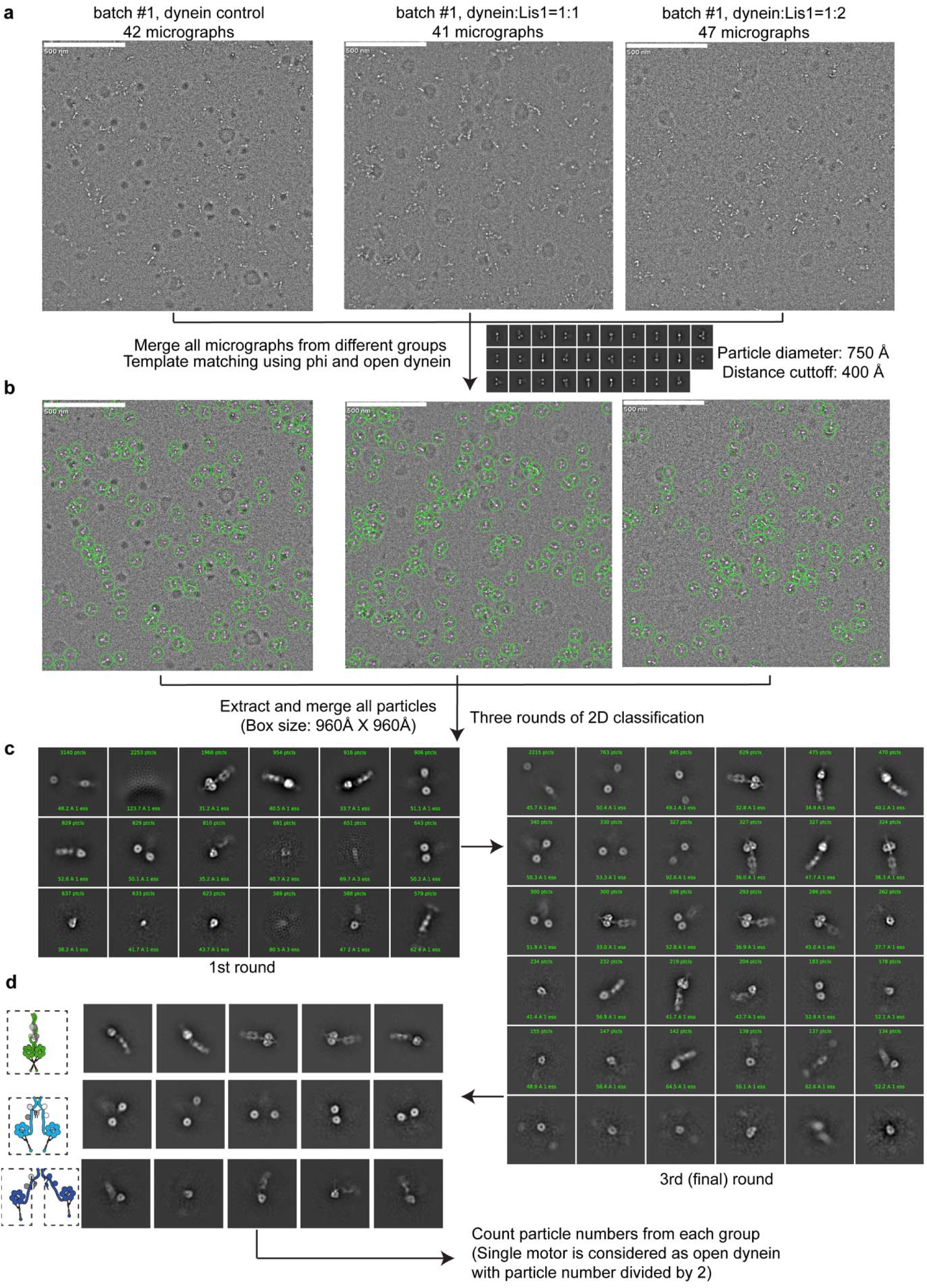
Workflow for negative-stain EM data processing. **a**, Representative micrographs for dynein alone (42 micrographs), dynein-Lis1 at 1:1 (41 micrographs), and dynein-Lis1 at 1:2 (47 micrographs) molar ratios from batch #1. **b**, Particle picking from representative micrographs in each dataset using a template matching approach based on Phi and open dynein models (particle diameter: 750 Å, distance cutoff: 400 Å). **c**, Three rounds of 2D classification were performed after extracting all particles (box size: 960 Å × 960 Å), yielding class averages of Phi dynein, open dynein motors, single motors, and junk particles. **d**, Final classified 2D averages showing Phi dynein, two-motor open dynein, and single-motor open dynein. The particle numbers for each group were counted, and single motors were considered as open dynein by dividing the total number of particles by two.

**Extended Data Fig. 4.**
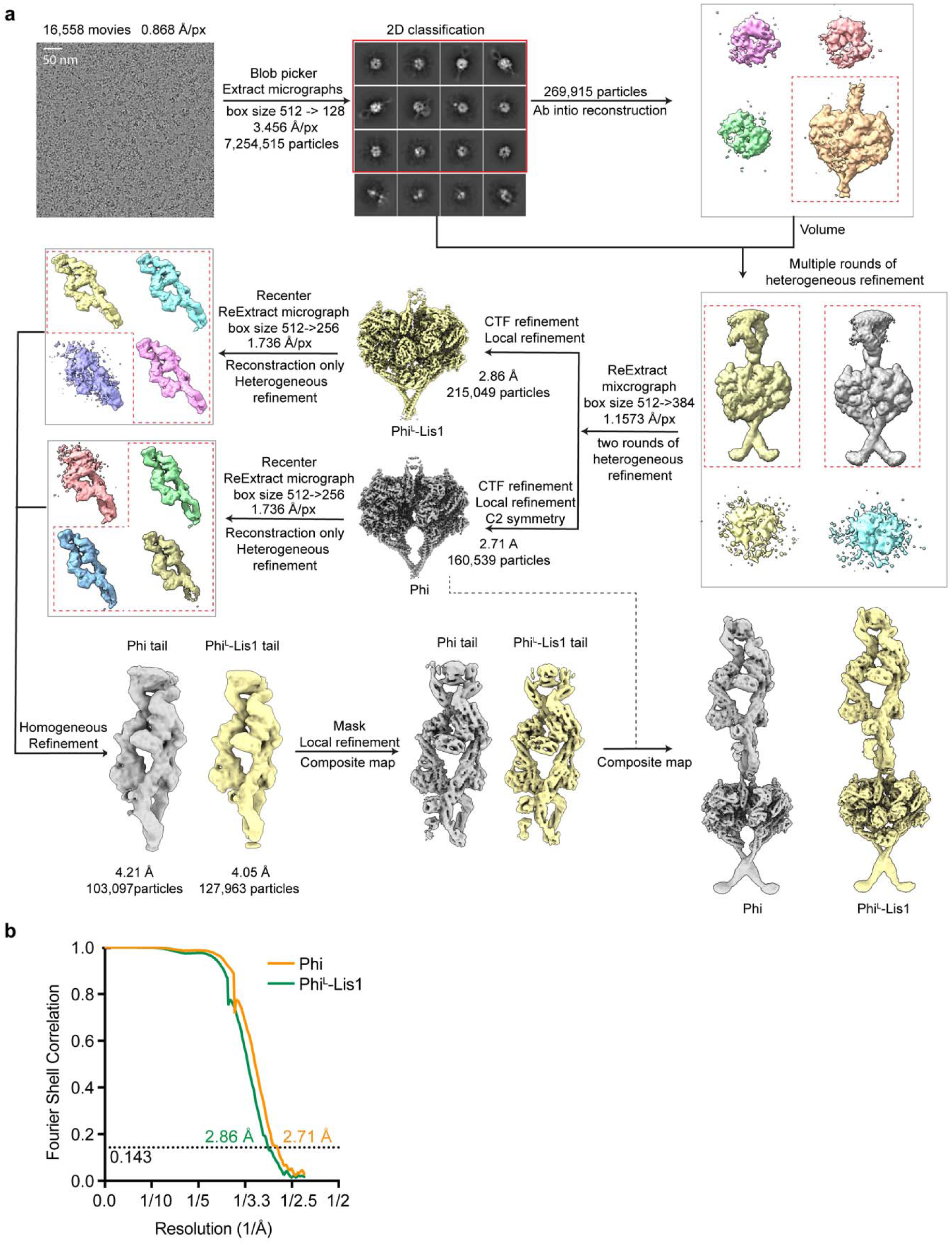
Cryo-EM data processing for the dynein-Lis1 dataset. **a**, A representative cryo-EM micrograph and the flowchart of cryo-EM data processing. **b**, Fourier Shell Correlation (FSC) curves showing the final resolution estimates for the motor domains of the Phi (2.71 Å) and Phi^L^-Lis1(2.86 Å) datasets.

**Extended Data Fig. 5.**
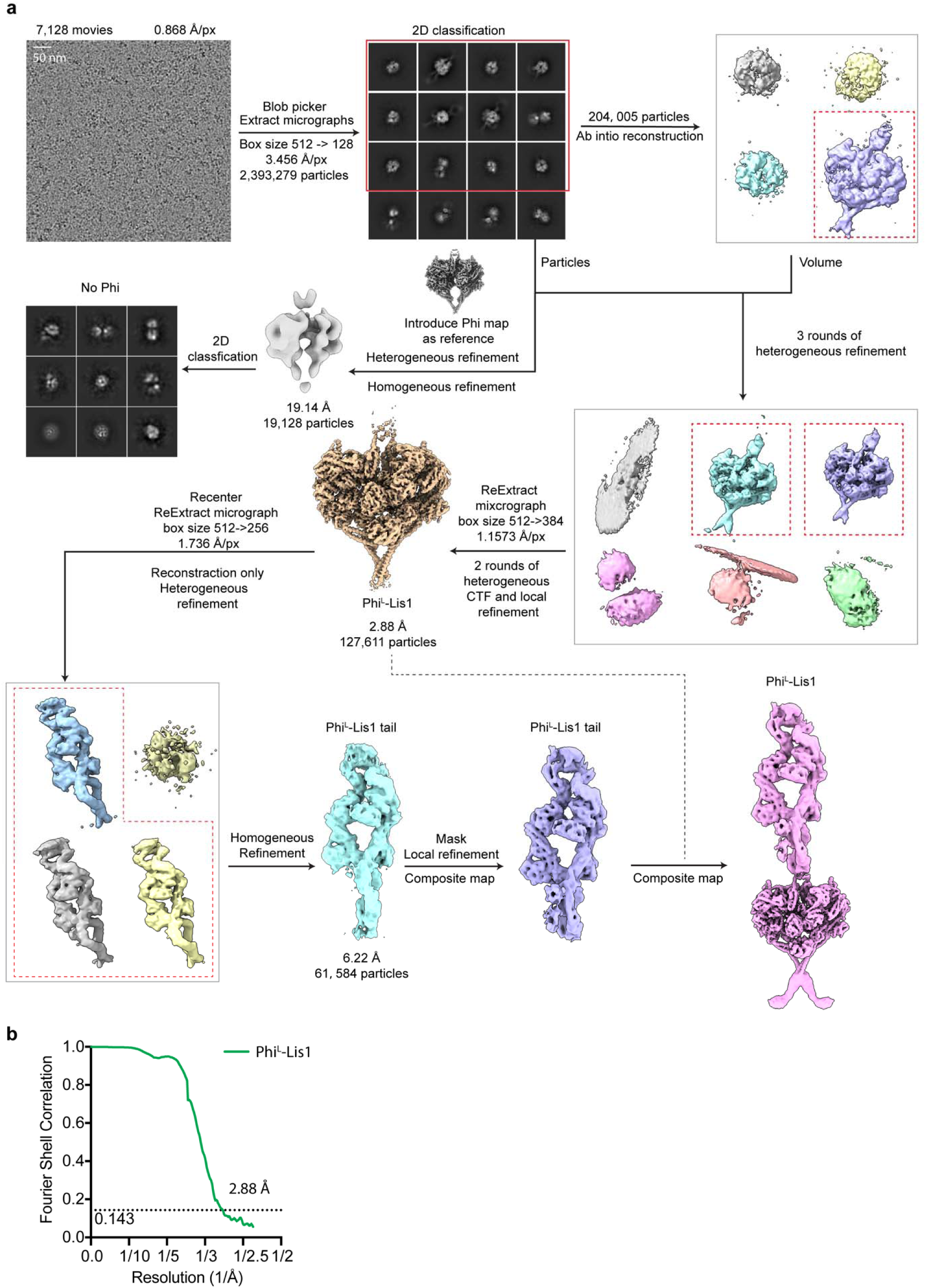
Cryo-EM data processing for the Nde1-Lis1-dynein dataset. **a**, A representative cryo-EM micrograph and the flowchart of cryo-EM data processing. **b**, Fourier Shell Correlation (FSC) curve showing the final resolution estimate for the motor domains of the Phi^L^-Lis1 (2.88 Å) dataset.

**Extended Data Fig. 6.**
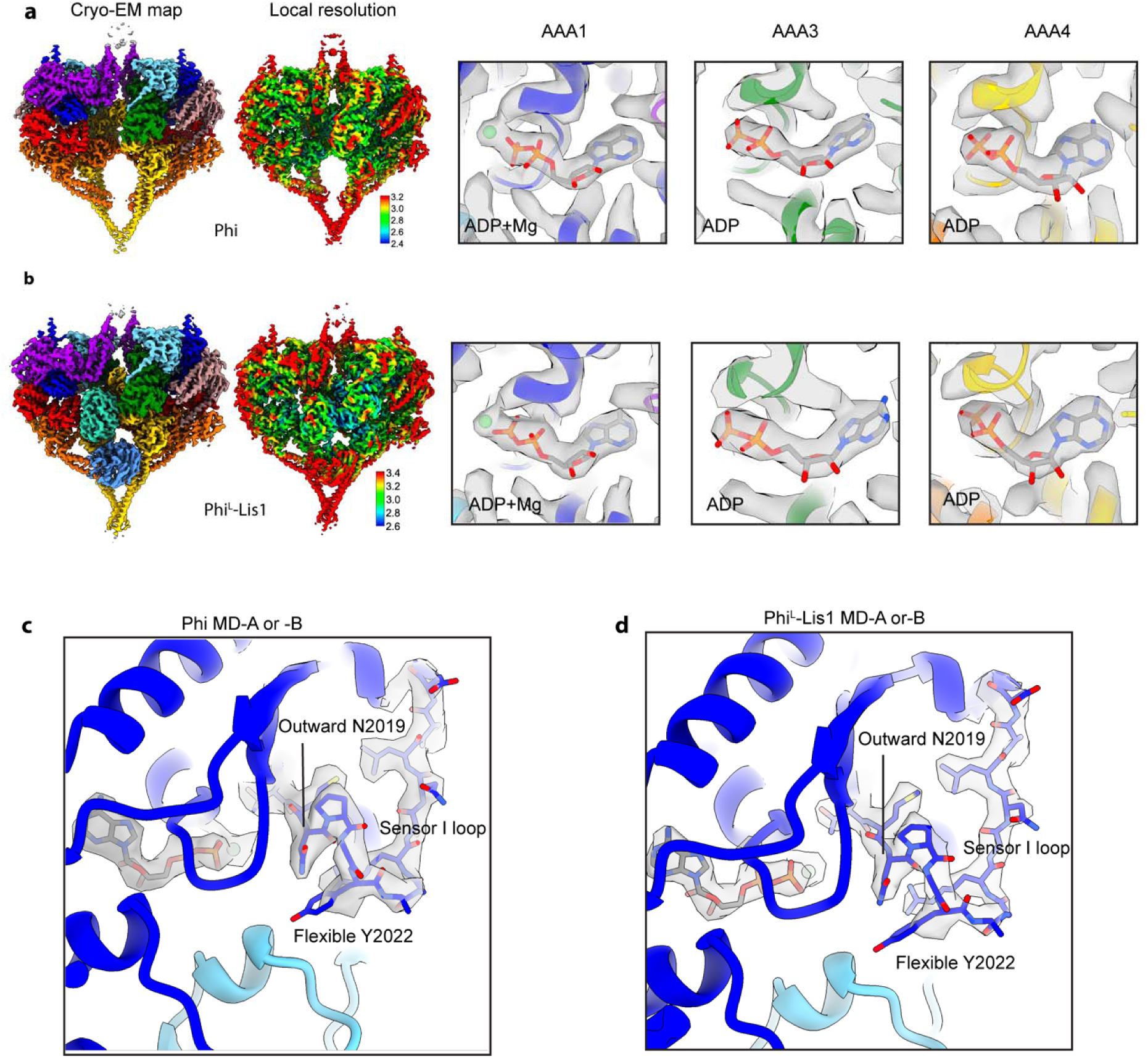
Comparison of local resolution, nucleotide binding in AAA1, AAA3, AAA4, and sensor-I loop conformation in MD-A of the Phi and Phi^L^-Lis1. Local resolution, and nucleotide binding states in MD-A at AAA1, AAA3, and AAA4 of the Phi (**a**) and Phi^L^-Lis1 (**b**). MD-A and -B share the same nucleotide binding in AAA1, AAA3, and AAA4 across both the Phi and Phi^L^-Lis1. The sensor-I loop adopts almost the same conformation in MD-A (or -B) of both Phi (**c**) and Phi^L^-Lis1 (**d**), indicating that Lis1 binding does not affect phosphate release. The color scheme is the same with Fig. 4.

**Extended Data Fig. 7.**
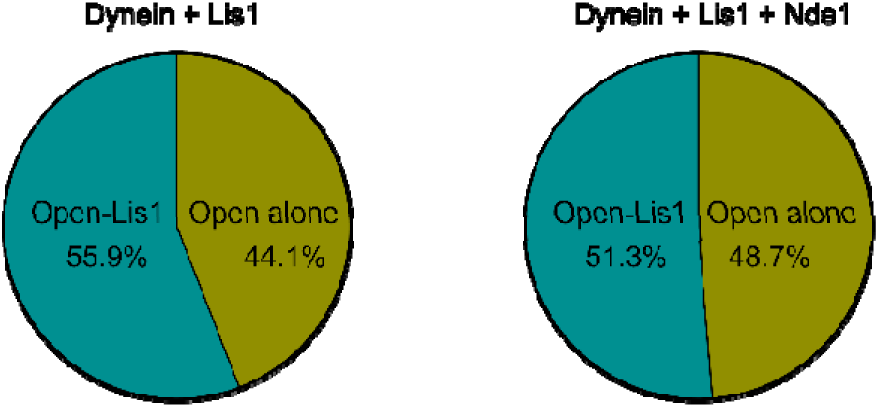
Comparison of Nde1’s effect on Lis1 binding to the open dynein. The particle numbers for open dynein-Lis1 and open dynein alone were quantified in both the dynein-Lis1 and dynein-Lis1-Nde1 datasets. These results indicate that Nde1 does not promote Lis1 binding to the open dynein motor.

**Extended Data Fig. 8.**
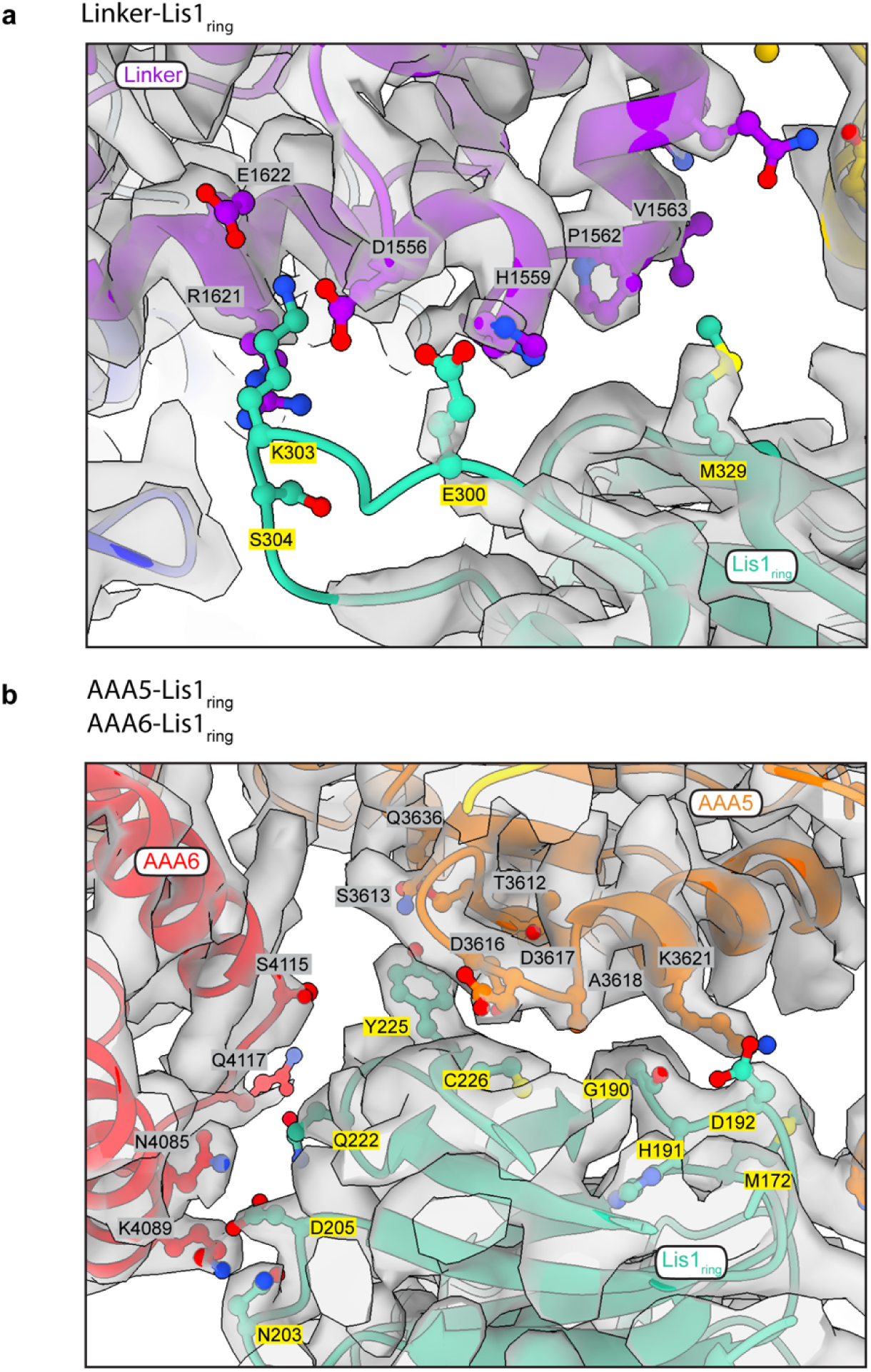
Density quality at the dynein MD-A and Lis1 interface in Phi^L^-Lis1. **a**, Flexible density at the linker-Lis1_ring_ interface, indicating dynamic interactions in this region. **b**, Well-defined density at the AAA5-Lis1_ring_ and AAA6-Lis1_ring_ regions, showing compact and stable interactions. The color scheme for the motor domains is consistent with Fig. 4.

**Extended Data Fig. 9.**
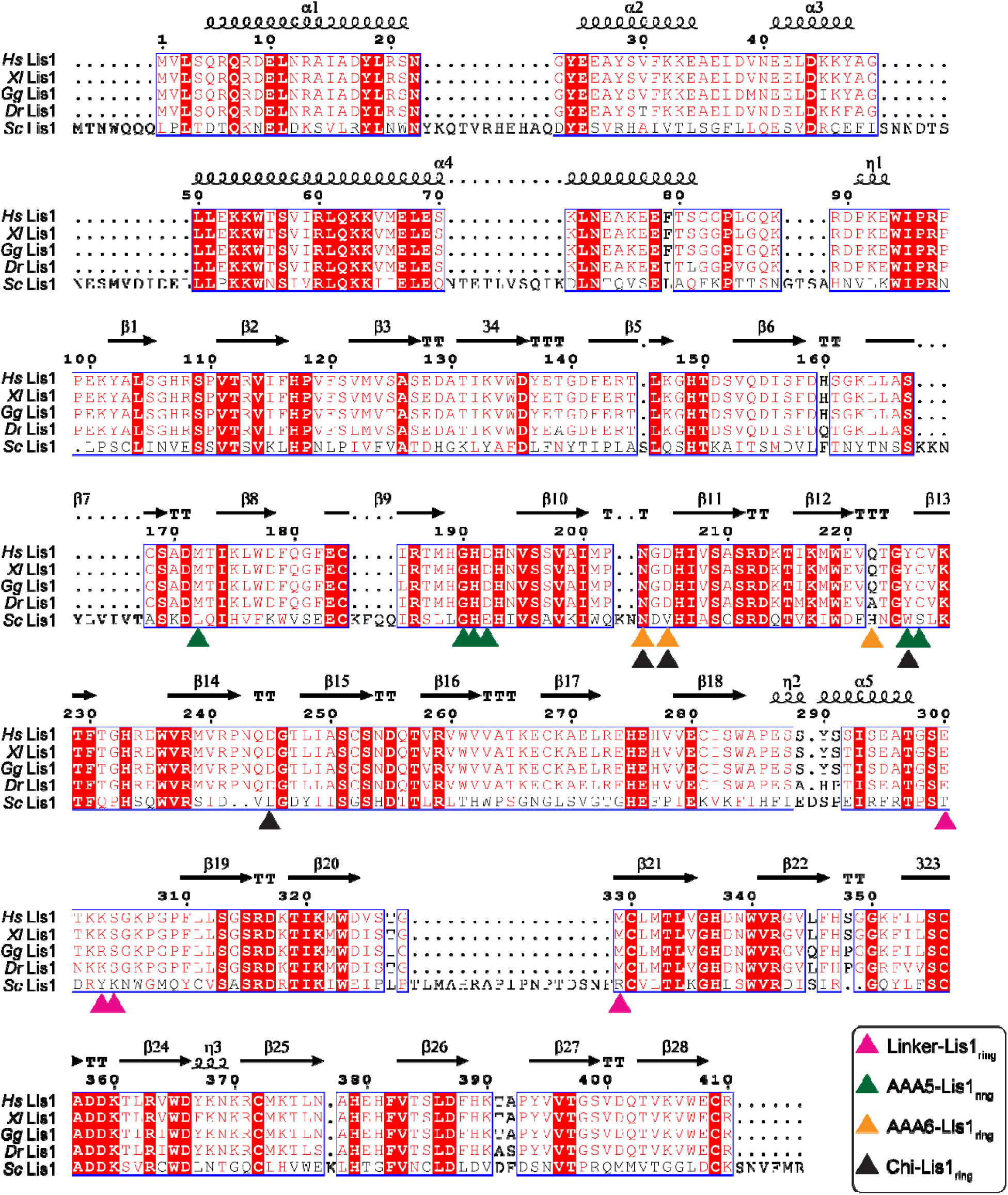
Sequence alignment of Lis1 homologs among multiple species. Sequence alignment of Lis1 proteins among *Homo sapiens* (Hs Lis1), *Xenopus laevis* (*Xl* Lis1), *Gallus gallus* (*Gg* Lis1), *Danio rerio* (*Dr* Lis1), and *Saccharomyces cerevisiae* (*Sc* Lis1). The secondary structure elements are placed on the top of the alignment. Strictly conserved residues are highlighted in shaded red boxes, while less-conserved residues are shown in open red boxes. In these open red boxes, red font indicates residues with similar polarity and high conservation, whereas black font represents residues with low similarity. The colored triangles represent key residues involved in interactions at the linker-Lis1_ring_ (purple), AAA5-Lis1_ring_ (green), and AAA6-Lis1_ring_ (orange) interfaces in Phi^L^-Lis1. The black triangle represents reported interactions at MD-A and Lis1 interface of modeled human Chi-Lis1 based on the yeast Chi-Lis1^37^. *S. cerevisiae* Lis1 shows more variation compared to the vertebrate species, suggesting greater evolutionary divergence.

**Extended Data Fig. 10.**
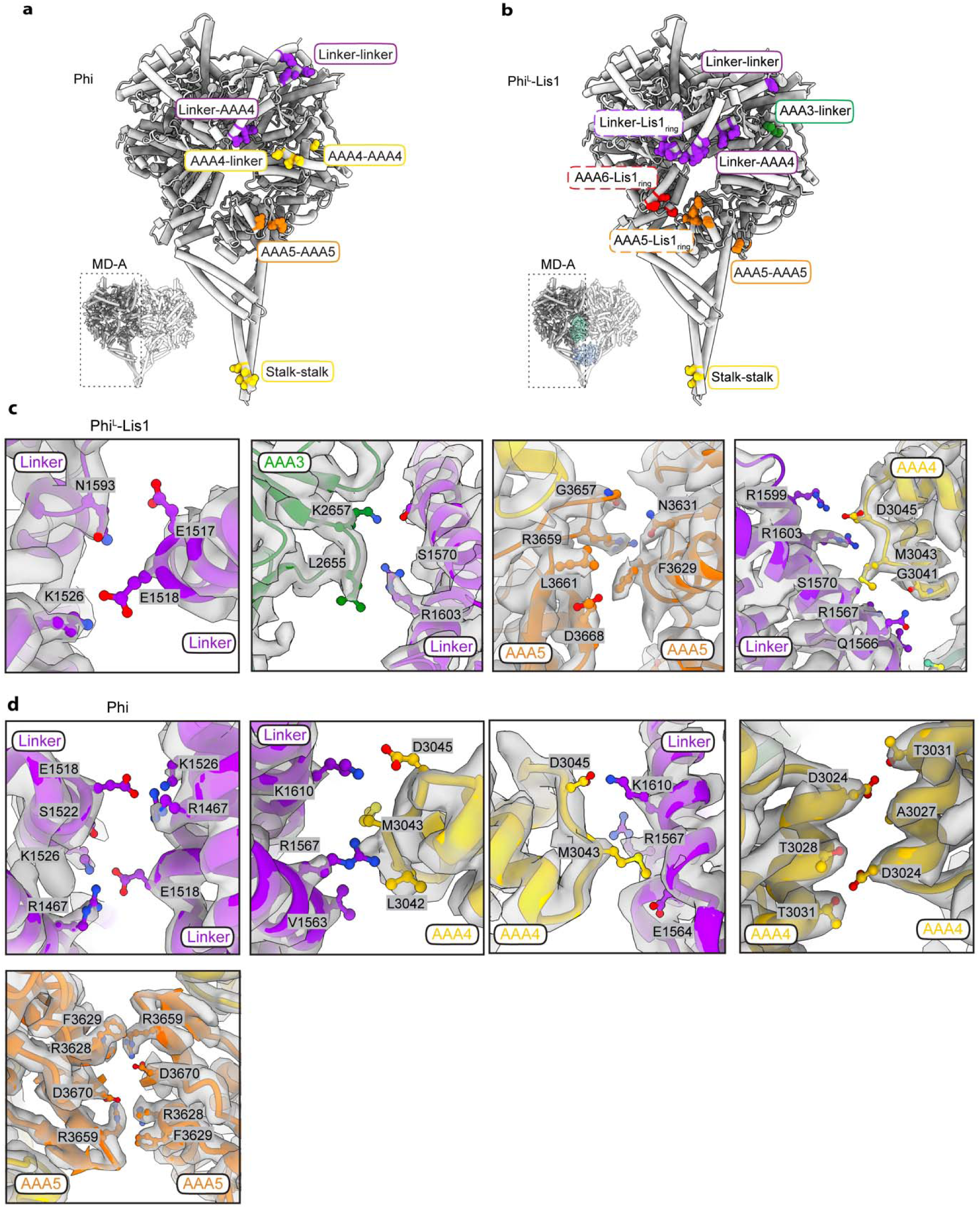
Comparison of motor domain A-B interfaces in Phi and Phi^L^-Lis1. **a**. Representative residues located on MD-A that are involved in the MD-A and MD-B interface of Phi, showing interactions at the linker-linker, linker-AAA4, AAA4-linker, AAA4-AAA4, AAA5-AAA5, and stalk-stalk interfaces. **b**, Representative residues located on MD-A of the Phi^L^-Lis1, involved in the MD-A and Lis1_ring_ interface (including linker-Lis1_ring_, AAA6-Lis1_ring_, and AAA5-Lis1_ring_), and the motor domain A-B interface (including linker-linker, AAA3-linker, linker-AAA4, AAA5-AAA5, and stalk-stalk interfaces). Residues in both panels (a) and (b) are displayed in sphere mode. **c**, Detailed view of the motor domain A-B interface in Phi^L^-Lis1, showing key residues involved in interactions at the linker-linker, AAA3-linker, AAA5-AAA5, and linker-AAA4 interfaces. **d**, Detailed view of motor domain A-B interface of Phi, showing key residues at the linker-linker, linker-AAA4, AAA4-linker, AAA4-AAA4, and AAA5-AAA5 interfaces.

**Extended Data Fig. 11.**
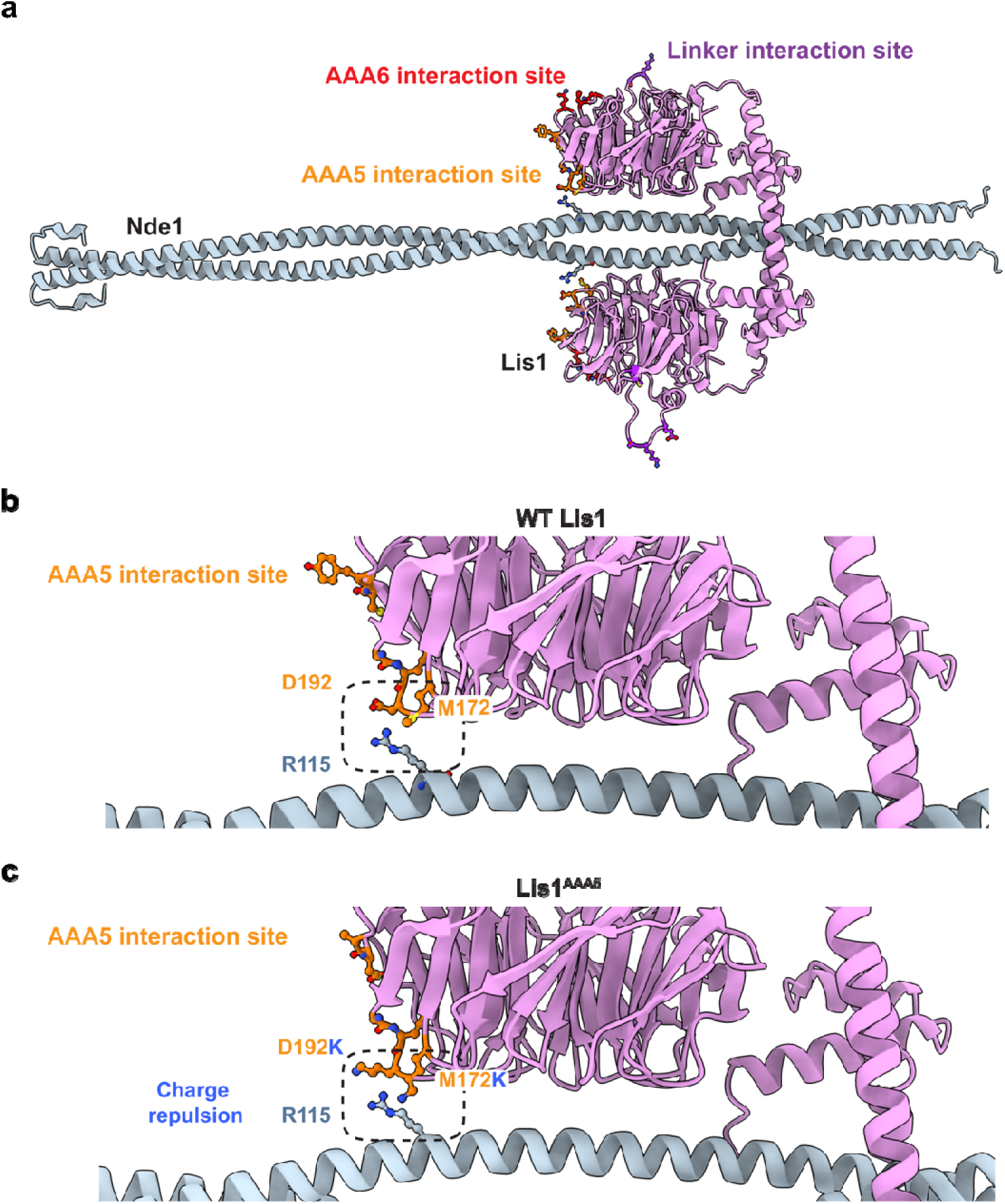
Nde1-Lis1 interface predicted by Alphafold. **a,** Predicted structure of the Nde1-Lis1 complex. Interactions involved in the Phi^L^ MD-A and Lis1 interface are shown on the Lis1 surface. **b**, D192 and M172 involved in AAA5-Lis1 interface also show contact with R115 of Nde1. **c**, D192K and M172K mutation of Lis1^AAA5^ show charge repulsion with R115 of Nde1. The prediction supports an overlap of the interfaces between the AAA5-Lis1 and Nde1-Lis1.

**Extended Data Fig. 12.**
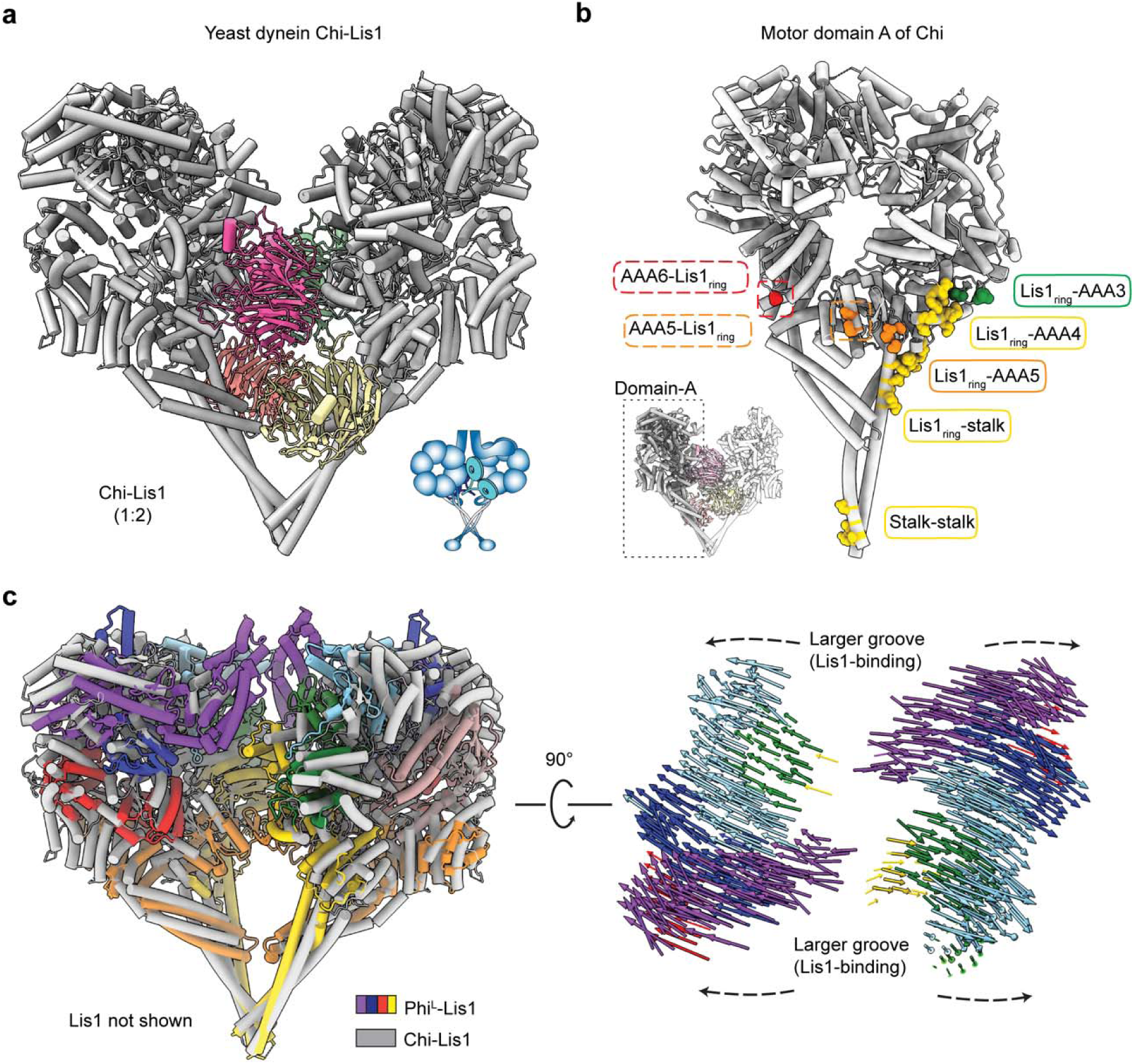
Comparison of yeast Chi-Lis1 and human Phi^L^-Lis1 motor domains. **a**, The structure of yeast Chi-Lis1 (PDB:8DZZ)^37^, showing two tail-truncated yeast dynein motor domains (grey) bound to two Lis1 dimers (colored, Chi-Lis1 1:2). **b**, Residues of MD-A that interact with Lis1_ring_ are located in AAA6-Lis1_ring_ and AAA5-Lis1_ring_ regions and highlighted with dashed rectangle. Representative residues of MD-A involved in the canonical Lis1_ring_ binding sites are located in Lis1_ring_-AAA3, Lis1_ring_-AAA4, Lis1_ring_-AAA5 and Lis1_ring_-stalk region. Interactions between MD-A and MD-B are in stalk-stalk region. Residues are displayed in sphere mode and are colored according to the subdomains in Fig. 4. **c**, Superimposition of the human Phi^L^-Lis1 and yeast Chi-Lis1 structures, showing that Chi-Lis1 adopts a more expanded conformation, with larger grooves on both the front and back sides compared to the more compact Phi^L^-Lis1 structure. Lis1 is hidden for clarity. Vectors represent interatomic distances of pairwise Cα atoms between the Phi^L^-Lis1 and Chi-Lis1 structures.

## Notes

### Competing Interest Statement

The authors have declared no competing interest.

### Summary of Updates

The PDB and EMDB IDs were removed from the manuscript.

## Reference

1. Reck-Peterson, S.L., Redwine, W.B., Vale, R.D. & Carter, A.P. The cytoplasmic dynein transport machinery and its many cargoes. Nat Rev Mol Cell Biol 19, 382–398 (2018).

2. McNally, F.J. Mechanisms of spindle positioning. J Cell Biol 200, 131–40 (2013).

3. Markus, S.M., Marzo, M.G. & McKenney, R.J. New insights into the mechanism of dynein motor regulation by lissencephaly-1. Elife 9(2020).

4. Willemsen, M.H. et al. Mutations in DYNC1H1 cause severe intellectual disability with neuronal migration defects. J Med Genet 49, 179–83 (2012).

5. Scoto, M. et al. Novel mutations expand the clinical spectrum of DYNC1H1-associated spinal muscular atrophy. Neurology 84, 668–679 (2015).

6. Poirier, K. et al. Mutations in TUBG1, DYNC1H1, KIF5C and KIF2A cause malformations of cortical development and microcephaly. Nat Genet 45, 639–47 (2013).

7. Guven, A., Gunduz, A., Bozoglu, T.M., Yalcinkaya, C. & Tolun, A. Novel NDE1 homozygous mutation resulting in microhydranencephaly and not microlyssencephaly. Neurogenetics 13, 189–94 (2012).

8. Lipka, J., Kuijpers, M., Jaworski, J. & Hoogenraad, C.C. Mutations in cytoplasmic dynein and its regulators cause malformations of cortical development and neurodegenerative diseases. Biochemical Society Transactions 41, 1605–1612 (2013).

9. Eschbach, J. & Dupuis, L. Cytoplasmic dynein in neurodegeneration. Pharmacol Ther 130, 348–63 (2011).

10. Urnavicius, L. et al. Cryo-EM shows how dynactin recruits two dyneins for faster movement. Nature 554, 202–206 (2018).

11. Singh, K. et al. Molecular mechanism of dynein-dynactin complex assembly by LIS1. Science 383, eadk8544 (2024).

12. Okada, K. et al. Conserved roles for the dynein intermediate chain and Ndel1 in assembly and activation of dynein. Nature Communications 14(2023).

13. Chaaban, S. & Carter, A.P. Structure of dynein-dynactin on microtubules shows tandem adaptor binding. Nature (2022).

14. Zhang, K. et al. Cryo-EM Reveals How Human Cytoplasmic Dynein Is Auto-inhibited and Activated. Cell 169, 1303–1314 e18 (2017).

15. Canty, J.T. & Yildiz, A. Activation and Regulation of Cytoplasmic Dynein. Trends in Biochemical Sciences 45, 440–453 (2020).

16. Neuwald, A.F., Aravind, L., Spouge, J.L. & Koonin, E.V. AAA+: A class of chaperone-like ATPases associated with the assembly, operation, and disassembly of protein complexes. Genome Res 9, 27–43 (1999).

17. Carter, A.P., Cho, C., Jin, L. & Vale, R.D. Crystal Structure of the Dynein Motor Domain. Science 331, 1159–1165 (2011).

18. Canty, J.T., Tan, R., Kusakci, E., Fernandes, J. & Yildiz, A. Structure and Mechanics of Dynein Motors. Annu Rev Biophys 50, 549–574 (2021).

19. Chai, P., et al. Cryo-EM Reveals the Mechanochemical Cycle of Reactive Full-length Human Dynein-1. BioRixv (2024).

20. Cianfrocco, M.A., DeSantis, M.E., Leschziner, A.E. & Reck-Peterson, S.L. Mechanism and regulation of cytoplasmic dynein. Annu Rev Cell Dev Biol 31, 83–108 (2015).

21. Amos, L.A. Brain dynein crossbridges microtubules into bundles. J Cell Sci 93 (Pt 1), 19–28 (1989).

22. Torisawa, T. et al. Autoinhibition and cooperative activation mechanisms of cytoplasmic dynein. Nature Cell Biology 16, 1118-+ (2014).

23. Garrott, S.R., Gillies, J.P. & DeSantis, M.E. Nde1 and Ndel1: Outstanding Mysteries in Dynein-Mediated Transport. Frontiers in Cell and Developmental Biology 10(2022).

24. Bradshaw, N.J., Hennah, W. & Soares, D.C. NDE1 and NDEL1: twin neurodevelopmental proteins with similar ‘nature’ but different ‘nurture’. Biomol Concepts 4, 447–64 (2013).

25. Monda, J.K. & Cheeseman, I.M. Nde1 promotes diverse dynein functions through differential interactions and exhibits an isoform-specific proteasome association. Molecular Biology of the Cell 29, 2336–2345 (2018).

26. Zhao, Y., Oten, S. & Yildiz, A. Nde1 promotes Lis1-mediated activation of dynein. Nat Commun 14, 7221 (2023).

27. Shu, T.Z. et al. Ndel1 operates in a common pathway with LIS1 and cytoplasmic dynein to regulate cortical neuronal positioning. Neuron 44, 263–277 (2004).

28. Stehman, S.A., Chen, Y., McKenney, R.J. & Vallee, R.B. NudE and NudEL are required for mitotic progression and are involved in dynein recruitment to kinetochores. Journal of Cell Biology 178, 583–594 (2007).

29. Wang, S.S. et al. Nudel/NudE and Lis1 promote dynein and dynactin interaction in the context of spindle morphogenesis. Molecular Biology of the Cell 24, 3522–3533 (2013).

30. McKenney, R.J., Vershinin, M., Kunwar, A., Vallee, R.B. & Gross, S.P. LIS1 and NudE Induce a Persistent Dynein Force-Producing State. Cell 141, 304–314 (2010).

31. Reiner, O. & Sapir, T. LIS1 functions in normal development and disease. Current Opinion in Neurobiology 23, 951–956 (2013).

32. Huang, J., Roberts, A.J., Leschziner, A.E. & Reck-Peterson, S.L. Lis1 Acts as a “Clutch” between the ATPase and Microtubule-Binding Domains of the Dynein Motor. Cell 150, 975–986 (2012).

33. Qiu, R.D., Zhang, J. & Xiang, X. LIS1 regulates cargo-adapter-mediated activation of dynein by overcoming its autoinhibition in vivo. Journal of Cell Biology 218, 3630–3646 (2019).

34. Elshenawy, M.M. et al. Lis1 activates dynein motility by modulating its pairing with dynactin. Nat Cell Biology 22, 570–578 (2020).

35. Htet, Z.M. et al. LIS1 promotes the formation of activated cytoplasmic dynein-1 complexes. Nature Cell Biology 22, 518-+ (2020).

36. McKenney, R.J. LIS1 cracks open dynein. Nature Cell Biology 22, 515–517 (2020).

37. Karasmanis, E.P. et al. Lis1 relieves cytoplasmic dynein-1 autoinhibition by acting as a molecular wedge. Nat Struct Mol Biol 30, 1357–1364 (2023).

38. Ton, W.D. et al. Microtubule-binding-induced allostery triggers LIS1 dissociation from dynein prior to cargo transport. Nature Structural & Molecular Biology (2023).

39. Kusakci, E. et al. Lis1 slows force-induced detachment of cytoplasmic dynein from microtubules. Nature Chemical Biology 20(2024).

40. Neer, E.J., Schmidt, C.J., Nambudripad, R. & Smith, T.F. The ancient regulatory-protein family of WD-repeat proteins. Nature 371, 297–300 (1994).

41. Emes, R.D. & Ponting, C.P. A new sequence motif linking lissencephaly, Treacher Collins and oral-facial-digital type 1 syndromes, microtubule dynamics and cell migration. Hum Mol Genet 10, 2813–20 (2001).

42. Mateja, A., Cierpicki, T., Paduch, M., Derewenda, Z.S. & Otlewski, J. The dimerization mechanism of LIS1 and its implication for proteins containing the LisH motif. Journal of Molecular Biology 357, 621–631 (2006).

43. Reimer, J.M., DeSantis, M.E., Reck-Peterson, S.L. & Leschziner, A.E. Structures of human dynein in complex with the lissencephaly 1 protein, LIS1. Elife 12(2023).

44. Marzo, M.G., Griswold, J.M. & Markus, S.M. Pac1/LIS1 stabilizes an uninhibited conformation of dynein to coordinate its localization and activity. Nature Cell Biology 22, 559-+ (2020).

45. Geohring, I.C. et al. A nucleotide code governs Lis1’s ability to relieve dynein autoinhibition. bioRxiv (2024).

46. Wynne, C.L. & Vallee, R.B. Cdk1 phosphorylation of the dynein adapter Nde1 controls cargo binding from G2 to anaphase. Journal of Cell Biology 217, 3019–3029 (2018).

47. Lam, C., Vergnolle, M.A., Thorpe, L., Woodman, P.G. & Allan, V.J. Functional interplay between LIS1, NDE1 and NDEL1 in dynein-dependent organelle positioning. J Cell Sci 123, 202–12 (2010).

48. Zhang, Y.F. et al. Nde1 is a Rab9 effector for loading late endosomes to cytoplasmic dynein motor complex. Structure 30, 386-+ (2022).

49. Doobin, D.J., Helmer, P., Carabalona, A., Bertipaglia, C. & Vallee, R.B. The Role of Nde1 phosphorylation in interkinetic nuclear migration and neural migration during cortical development. Mol Biol Cell 35, ar129 (2024).

50. Pei, Z. et al. The Expression and Roles of Nde1 and Ndel1 in the Adult Mammalian Central Nervous System. Neuroscience 271, 119–136 (2014).

51. Derewenda, U. et al. The structure of the coiled-coil domain of Ndel1 and the basis of its interaction with Lis1, the causal protein of Miller-Dieker lissencephaly. Structure 15, 1467–81 (2007).

52. Garrott, S.R. et al. Ndel1 disfavors dynein-dynactin-adaptor complex formation in two distinct ways. J Biol Chem 299, 104735 (2023).

53. Ye, F. et al. DISC1 Regulates Neurogenesis via Modulating Kinetochore Attachment of Ndel1/Nde1 during Mitosis. Neuron 96, 1204 (2017).

54. McKenney, R.J., Weil, S.J., Scherer, J. & Vallee, R.B. Mutually Exclusive Cytoplasmic Dynein Regulation by NudE-Lis1 and Dynactin. Journal of Biological Chemistry 286, 39615–39622 (2011).

55. Nyarko, A., Song, Y.J. & Barbar, E. Intrinsic Disorder in Dynein Intermediate Chain Modulates Its Interactions with NudE and Dynactin. Journal of Biological Chemistry 287, 24884–24893 (2012).

56. Moon, H.M. et al. LIS1 controls mitosis and mitotic spindle organization via the LIS1-NDEL1-dynein complex. Hum Mol Genet 23, 449–66 (2014).

57. Zylkiewicz, E. et al. The N-terminal coiled-coil of Ndel1 is a regulated scaffold that recruits LIS1 to dynein. J Cell Biol 192, 433–45 (2011).

58. Wang, S.S. & Zheng, Y.X. Identification of a novel dynein binding domain in nudel essential for spindle pole organization in Xenopus egg extract. Journal of Biological Chemistry 286, 587–593 (2011).

59. Efimov, V.P. Roles of NUDE and NUDF Proteins of Aspergillus nidulans: Insights from Intracellular Localization and Overexpression Effects. Molecular Biology of the Cell 14, 871–888 (2003).

60. Toropova, K. et al. Lis1 regulates dynein by sterically blocking its mechanochemical cycle. Elife 3(2014).

61. Cianfrocco, M.A. et al. Lis1 has Two Distinct Modes of Regulating Dynein’s Mechanochemical Cycle. Biophysical Journal 112, 43a–43a (2017).

62. Gillies, J.P. et al. Structural basis for cytoplasmic dynein-1 regulation by Lis1. Elife 11(2022).

63. Schlager, M.A., Hoang, H.T., Urnavicius, L., Bullock, S.L. & Carter, A.P. In vitro reconstitution of a highly processive recombinant human dynein complex. EMBO J 33, 1855–68 (2014).

64. Urnavicius, L. et al. The structure of the dynactin complex and its interaction with dynein. Science 347, 1441–1446 (2015).

65. Casanal, A., Lohkamp, B. & Emsley, P. Current developments in Coot for macromolecular model building of Electron Cryo-microscopy and Crystallographic Data. Protein Sci 29, 1069–1078 (2020).

66. Kidmose, R.T. et al. Namdinator - automatic molecular dynamics flexible fitting of structural models into cryo-EM and crystallography experimental maps. IUCrJ 6, 526–531 (2019).

67. Afonine, P.V. et al. Real-space refinement in PHENIX for cryo-EM and crystallography. Acta Crystallogr D Struct Biol 74, 531–544 (2018).

